# Reactivity-based screening for citrulline-containing natural products reveals a family of bacterial peptidyl arginine deiminases

**DOI:** 10.1101/2020.07.16.207027

**Authors:** Lonnie A. Harris, Patricia M. B. Saint-Vincent, Xiaorui Guo, Graham A. Hudson, Douglas A. Mitchell

**Author notes:** Correspondence, S. Mathews Avenue, Urbana, IL 61802.

## Abstract

Ribosomally synthesized and post-translationally modified peptides (RiPPs) are a family of natural products defined by a genetically encoded precursor peptide that is tailored by associated biosynthetic enzymes to form the mature product. Lasso peptides are a class of RiPP defined by an isopeptide linkage between the N-terminal amine and an internal Asp/Glu residue with the C-terminus threaded through the macrocycle. This unique lariat topology, which provides considerable stability towards heat and proteases, has stimulated interest in lasso peptides as potential therapeutics. Post-translational modifications beyond the class-defining, threaded macrolactam have been reported, including one example of arginine deimination to yield citrulline. Although a citrulline-containing lasso peptide (i.e., citrulassin) was serendipitously discovered during a genome-guided campaign, the gene(s) responsible for arginine deimination has remained unknown. Herein we describe the use of reactivity-based screening to discriminate bacteria that produce arginine-versus citrulline-bearing citrulassins, culminating in the discovery and characterization of 11 new lasso peptide variants. Phylogenetic profiling identified a distally encoded peptidyl arginine deiminase (PAD) gene ubiquitous to the citrulline-containing variants. Absence of this gene correlated strongly with citrulassin variants only containing arginine (*des*-citrulassin). Heterologous expression of the PAD in a non-citrulassin producer resulted in the production of the deiminated analog, confirming PAD involvement in arginine deimination. The family of PADs were then bioinformatically surveyed for a deeper understanding of its genomic context and potential role in post-translational modification of RiPPs.

## Introduction

Ribosomally synthesized and post-translationally modified peptides (RiPPs) are a family of natural products (NPs) defined by a common biosynthetic logic, wherein a genetically encoded precursor peptide is modified by associated biosynthetic enzymes to form the mature NP.^1, 2^ Precursor peptides are typically bipartite, composed of an N-terminal leader region responsible for enzyme recognition and a C-terminal core region which is modified and cleaved from the leader region to form the mature RiPP.^1, 2^ Because substrate recognition is guided by the leader sequence, RiPP biosynthetic enzymes are typically permissive of sequence changes in the core region, facilitating access to analogs.^3^ One RiPP class that has attracted interest are the lasso peptides, which are defined by a unique lariat topology that grants them thermal stability and protease resistance.^4^ These properties, in addition to the substrate permissiveness of several characterized lasso biosynthetic enzymes, have engendered interest in the design of lasso peptides therapeutics.^5, 6, 7, 8, 9^

After ribosomal synthesis of the lasso precursor peptide, the next biosynthetic step involves RiPP recognition element (RRE) engagement of the precursor peptide, which occurs through binding to a recognition sequence within the N-terminal leader region. The RRE domain mediates recruitment of the subsequent enzymes^10, 11^ and is found as either a discretely encoded protein or fused to the leader peptidase. Upon RRE binding, the leader peptidase removes the leader region from the precursor peptide.^12, 13^ The characteristic lariat topology is then formed by an ATP-dependent lasso cyclase, which installs a macrolactam linkage between the newly formed N-terminus of the core with the carboxylate of a downstream Asp or Glu acceptor residue, trapping the C-terminal tail within the ring.^12, 13^

As with other RiPP classes, post-translational modifications (PTMs) beyond the class-defining, threaded macrolactam are known, among them are disulfide formation (e.g. BI-32169 and others),^14^ C-terminal *O*-methylation (e.g. lassomycin),^15^ Asp β-hydroxylation (e.g. canucin A),^16^ Lys ε-acylation (e.g. albusnodin),^17^ Trp 7-hydroxylation (e.g. RES-701-2),^18^ epimerization to D-Trp (e.g. MS-271),^19^ Ser *O*-phosphorylation (e.g. paeninodin),^20^ and Arg deimination (e.g. citrulassin A).^21^ The latter modification was discovered during a genomics-guided discovery of novel lasso peptides, enabled by the bioinformatics tool Rapid ORF Description and Evaluation Online (RODEO). Briefly, RODEO uses profile hidden Markov models (pHMMs), heuristic scoring, and supervised machine learning to identify putative RiPP biosynthetic gene clusters (BGCs) and score potential precursors based on known family members.^21^ Since its initial application to lasso peptides, additional RODEO modules have been developed and used to mine and classify other RiPP classes, such as thiopeptides,^22^ lanthipeptides,^23^ sactipeptides, and ranthipeptides.^24^

During our previous RODEO-guided lasso peptide discovery effort, an uncharacterized family of 55 lasso peptides was bioinformatically identified (**Figure 1)**.^21^ The first member of this family, citrulassin A, was isolated and characterized from *Streptomyces albulus* NRRL B-3066. Citrulassin A was so named owing to an unprecedented citrulline at position nine of the core region (Cit9) rather than the genetically encoded Arg. Because citrulline is a non-proteinogenic amino acid, it was hypothesized that a peptidyl arginine deiminase (PAD) might be responsible for forming the citrulline moiety of citrulassin A, as this enzyme family carries out post-translational deimination of Arg in other biological contexts. However, genome sequencing of *S. albulus* NRRL B-3066 revealed no enzymes with predicted PAD activity within 15 open-reading frames of the citrulassin A cyclase (WP_079136914.1). Moreover, heterologous expression of the citrulassin A BGC with large flanking regions (∼20 kbp on each side of the BGC) in *Streptomyces lividans* resulted in the production of *des*-citrulassin A, which contains Arg at position nine of the core (Arg9) instead of citrulline (**Figure 1**). Given this result, the gene(s) responsible for Arg deimination was presumed to be distally encoded to the citrulassin A BGC.^21^

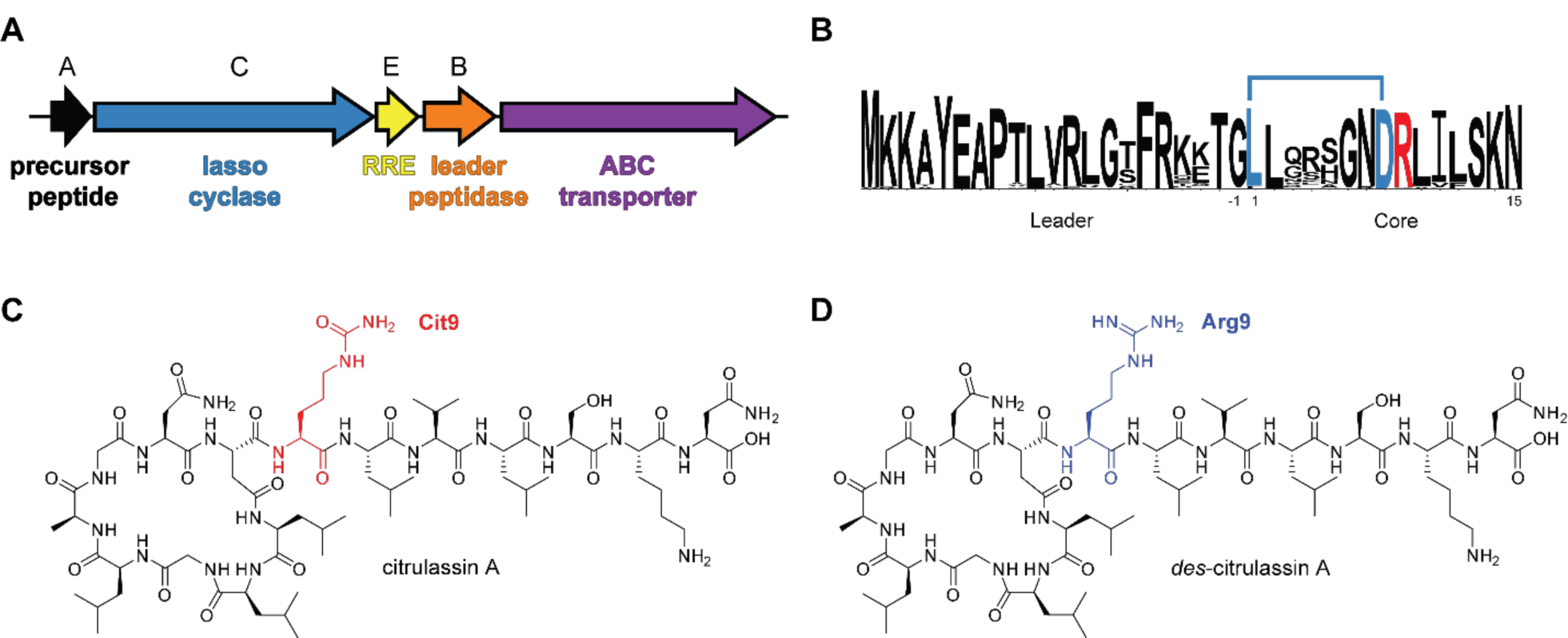
(A) General lasso peptide biosynthetic gene cluster architecture. (B) Multiple sequence alignment (http://weblogo.berkeley.edu/logo.cgi)^8^ of the 181 bioinformatically identified citrulassin precursor peptides. Isopeptide bond formation between the Asp side chain and N-terminus of the core region is indicated with a blue bar. The Arg residue modified to citrulline in citrulassin A is red. (C) Two-dimensional structure of citrulassin A from *S. albulus* NRRL B-3066, with Cit9 highlighted in red. (D) Two-dimensional structure of *des*-citrulassin A heterologously expressed in *S. lividans*, with Arg9 highlighted in blue. Non-threaded versions of the structures are shown for improved clarity.

To the best of our knowledge, only one bacterial PAD has been characterized. This PAD is from *Porphyromonas gingivalis*, conserved amongst periodontal pathogens,^25^ and contributes to *P. gingivalis* virulence. The *P. gingivalis* PAD enzyme deiminates human fibrinogen, leading to gingivitis and an increased risk of rheumatoid arthritis.^26^ This PAD virulence factor (protein family^27^ PF08527, PAD_porph) is distinct from human PADs (protein family PF03068), of which the PAD4 isoform is the best studied. PAD4 deiminates histones, which can affect transcription as well as cellular differentiation, and has been implicated in the onset of rheumatoid arthritis and in certain cancers.^28^

To identify the putative PAD involved in the deimination of citrulassin A, further screening of citrulassin producers for the presence of citrulline was required prior to pursuing a comparative genomic approach. Arg deimination, however, is difficult to discern by mass spectrometry, as the PTM results in a small mass deviation (replacement of NH with O, +0.98 Da) that is easily overlooked, especially when both Arg- and citrulline-containing congeners are produced. Moreover, deimination of Asn/Gln to the corresponding carboxylic acid is isobaric with deamidation, a commonly observed, spontaneous side chain modification, further complicating identification of deimination.^29^ Building off of precedent in natural product discovery and proteomics, we have previously described strategies for the rapid discovery of NPs bearing specific functional groups in the extracts of cultured bacteria, termed reactivity-based screening (RBS) (**Figure S1**).^30^ RBS involves the chemoselective labeling of an organic functional group, whereby NPs bearing the targeted functional group are rapidly detected via comparative mass spectrometry (MS) of reacted and unreacted samples. RBS has enabled the discovery and subsequent characterization of RiPPs, non-ribosomal peptides, polyketides, and hybrids thereof (e.g. cyclothiazomycin C,^31^ *deimino*-antipain,^32^ hygrobafilomycin JBIR-100,^33^ and tirandalydigin,^34^ *etc.*). RBS is therefore a valuable tool for bridging bioinformatics-guided NP identification with NP production. Additionally, an RBS-based NP discovery strategy permits rapid dereplication of already-characterized NPs,^31^ detection of low-abundance species using sensitive MS-based techniques such as matrix-assisted laser/desorption ionization time-of-flight MS (MALDI-TOF-MS), and can employ variable probe chemistries that enhance NP detection due to unique isotopic patterns of labeled compounds or selective enrichment through affinity purification.^32, 35, 36^

Herein, we describe the expansion of RBS as a tool to aid in the discovery of the PAD responsible for deimination activity on the citrulassin family of lasso peptides. Repurposing the work of Thompson,^37, 29^ a phenylglyoxal probe was employed to specifically label primary ureido groups over primary guanidino groups. This probe was used to survey for the presence of citrulline in a subset of lasso peptides and enable phylogenetic profiling, thus revealing a putative PAD in each citrulassin producer. A broader bioinformatic survey of bacterial PADs was also performed, revealing their taxonomic breadth and diversity of genomic contexts. Lastly, we heterologously express a PAD from *Streptomyces glaucescens* and demonstrate the deimination of a citrulassin precursor peptide *in vivo*, providing the first gene-to-function link for RiPP citrulline formation.

## Results and Discussion

### Validation of 3-bromophenylglyoxal as a primary ureido-selective probe

Previous work has demonstrated selective labeling of citrulline in eukaryotic cellular extracts at low pH with phenylglyoxal probes (**Figure 2**).^29, 37^ Under sufficiently acidic conditions, guanidino groups are protonated and thus unreactive towards the glyoxal moiety; however, ureido groups remain nucleophilic and thus react rapidly.^37^ The reaction of citrulline with phenylglyoxal has been further elaborated to readily identify labeled species in complex samples by employing brominated phenylglyoxal derivatives. The bromine provides a distinct isotopic pattern (∼1:1 ^79^Br/^81^Br) while the UV-absorbing arene increases signal intensity in MALDI-TOF-MS.^38^ Moreover, the probe can easily be modified to enrich for labeled compounds using a biotin-conjugated glyoxal probe.^27^ Thus, we imagined that 3-bromophenylglyoxal (**1**) could be deployed to discover primary ureido containing NPs from bacterial extracts. As predicted from the known reactivity of glyoxal, the free amino acid L-citrulline was robustly modified by **1** while L-Arg was unmodified under identical conditions (**Figure S2**). To evaluate the suitability of reaction within the context of a cellular extract, *S. albulus* NRRL B-3066 (the native producer of citrulassin A) and *Streptomyces lividans* 3H4 (a heterologous producer of *des*-citrulassin A)^20^ were grown for 7 d and metabolites were extracted from the cell-surface with methanol. When treated with **1** under identical reaction conditions, citrulassin A was nearly fully labeled while negligible product was observed for *des*-citrulassin A (**Figure 2**). Moreover, *deimino*-antipain, another citrulline-containing compound produced by *S. albulus* NRRL B-3066, was labeled under these conditions (**Figure S3**). Extract from *S. lividans* 3H4^21^ heterologously producing antipain confirmed selectivity for citrulline over Arg, with antipain displaying negligible labeling by **1**. Taken together, these experiments validate **1** as useful in the rapid identification of NPs containing a primary ureido group.

**Figure 2.**
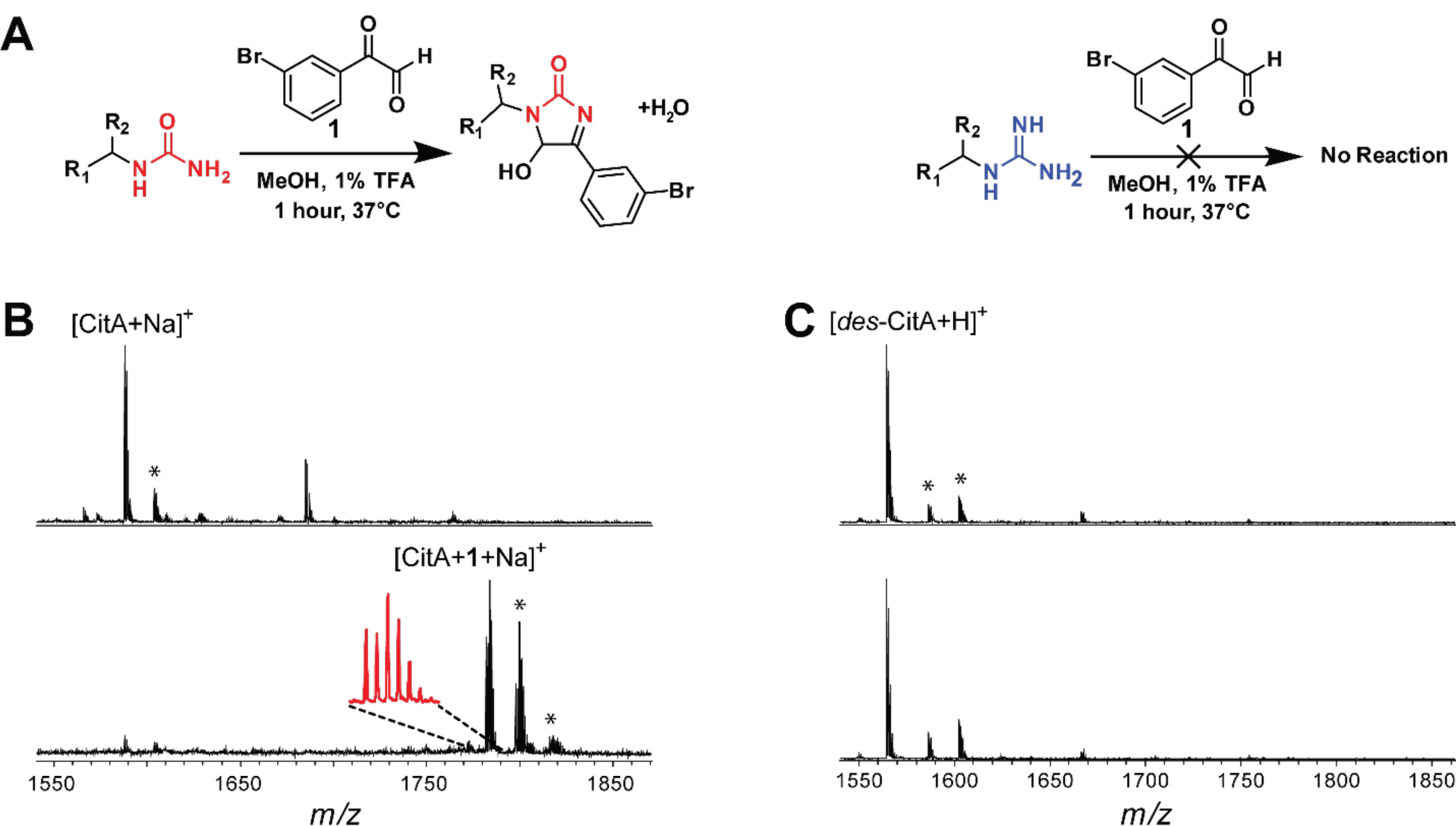
Validation of 3-bromophenylglyoxal (**1**) selectivity. (**A**) Reaction conditions that selectively label citrulassin (left) over *des*-citrulassin (right). TFA = trifluoroacetic acid. (**B**) MALDI-TOF mass spectra of a methanolic extract of *Streptomyces albulus* (citrulassin A producer) prior to addition of **1** (*top*) and after reaction with **1** (*bottom*). Magnified inset shows the ^79^Br:^81^Br isotopic pattern. Peaks indicated with an asterisk are (from left to right): [CitA+K]^+^, [CitA+**1**+K]^+^, and [CitA+**1**+K+H2O]^+^. (**C**) MALDI-TOF mass spectra of a methanolic extract of *des*-citrulassin A heterologously expressed in *Streptomyces lividans* (ref. 21) prior to addition of **1** (*top*) and after reaction with **1** (*bottom*). Peaks indicated with an asterisk are (from left to right): [*des*-CitA+Na]^+^ and [*des*-CitA+K]^+^.

### 3-bromophenylglyoxal (1) screening of citrulassin producers

With **1** validated, we next sought to identify which members of the citrulassin family of lasso peptides contain a post-translationally derived citrulline. Previous screening of bacterial extracts suggested that not all citrulassins contain citrulline (the Arg-containing counterpart is referred to as *des*-citrulassin),^21^ but owing to a small mass difference between guanido- and ureido-containing products, we tested all putative citrulassin producers through reactivity with **1**. To identify probable citrulassin producers, a BLAST-p search was performed using the citrulassin A cyclase as a query (WP_079136914.1). Cyclases with an e-value <10^−200^ were then subjected to RODEO analysis. The identified citrulassin precursor peptides were remarkably well conserved, with only a few positions in the ring and loop regions showing sequence variation (**Figure 1** and **Supplemental Dataset 1**). In addition, co-occurrence analysis indicated that no putative PADs are encoded in the genomic neighborhood of the citrulassin BGCs, in accord with the earlier heterologous expression study.^21^ With this list in hand, a selection of 109 strains predicted to contain citrulassin BGCs were obtained from the Agriculture Research Service (ARS) Culture Collection and grown for 10 d on a variety of media at 30 °C. Afterwards, the bacteria were harvested and metabolites extracted from the cell surface using MeOH for 2 h (see methods). Reactions with **1** were performed directly on the methanolic extracts and subsequently dried, reconstituted in methanol, and analyzed by MALDI-TOF-MS for the presence of primary ureido-containing metabolites. Extracts from which a predicted citrulassin were detected (n = 12) were screened for Arg deimination using reactivity towards **1** (**Table 1**). Extracts from six of these strains underwent labeling and were subjected to high-resolution and tandem mass spectrometry (HR-MS/MS), which confirmed the molecular formula and location of citrulline(s) (**Figure S5**). Several citrulassins identified contain Arg within the ring, typically at position four, in addition to the conserved Arg9 encoded within the loop region. Citrulline formation was confined to position 9 for all citrulassins with the exception of one variant from *Streptomyces* sp. NRRL S-920, which contained a second citrulline moiety within the ring. The extent of deimination was accurately reflected by reactivity towards **1**, with all singly deiminated citrulassins showing exactly one labeling event, while the doubly deiminated citrulassin was labeled twice (**Figure S5**). Thus, this approach is useful for determining the stoichiometry of targetable functional groups on a compound of interest.

**Table 1:**
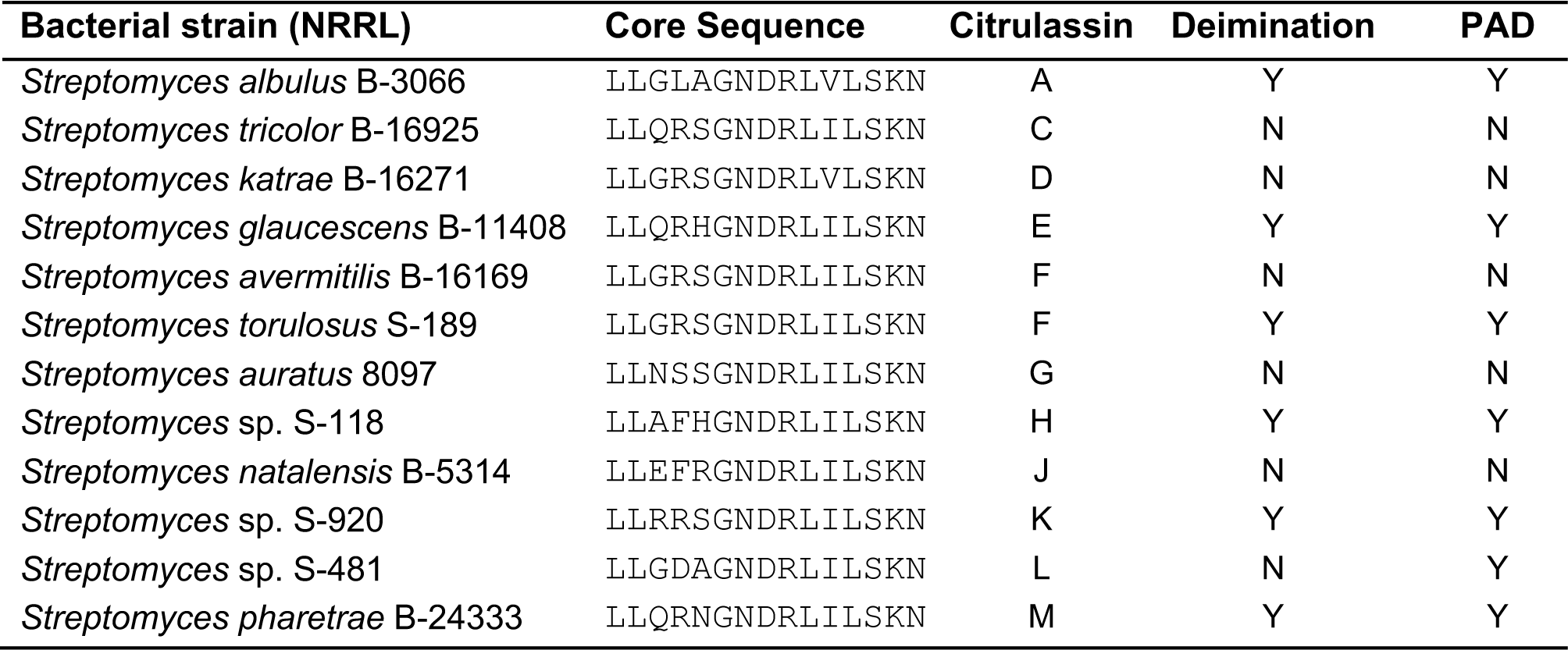
Characterized citrulassins.

### Phylogenetic profiling identifies a putative PAD for citrulline formation

Knowing which sequenced *Streptomyces* strains produce citrulassin versus *des*-citrulassin allowed a phylogenetic profiling strategy to identify the putative gene(s) responsible for Arg deimination. Using PhyloProfile,^39, 40^ a search was conducted for genes present in the *S. albulus* genome that were also present in four other citrulassin producers but not in four *des*-citrulassin producers. This analysis returned 32 *S. albulus* genes that correlated with the citrulline phenotype, one of which encoded a hypothetical protein containing a PAD domain (**Supplemental Dataset 1**). This gene encodes a member of protein family PF03068 and thus is homologous to human PADs. No significant sequence homology is shared between the *S. albulus* PAD (WP_064069847.1) and the only other characterized bacterial PAD from *P. gingivalis* (protein family PF04371, WP_005873463.1).^27,25^ (**Table S3)** Position-specific iterative BLAST (PSI-BLAST)^41^ of citrulassin producers revealed a correlation between the genome of a bacterial strain encoding a PAD and the presence of citrulline in the mature citrulassin, with only one strain producing *des*-citrulassin while having a PAD in the genome. Analogously, all organisms that lacked a PAD were associated with *des*-citrulassin productions. However, the small sample size prevented a statistically relevant correlation between genotype and chemotype from being established.^42^ Nonetheless, having identified a candidate PAD, we set out to further examine bacterial members of protein family PF03068.

### Bioinformatic survey of PF03068 bacterial PADs

While the *S. albulus* PAD is classified within the same protein family as eukaryotic PADs (PF03068), the sequence similarity is only 42-47% between each of the five human PAD isoforms and the PAD from *S. albulus* (**Table S4**). Despite this modest sequence similarity, an alignment of the *S. albulus* PAD with human PAD4, the best-characterized isoform, shows conservation of functionally important residues. The human PADs are calcium-dependent enzymes with a conserved catalytic Cys645 and stabilizing triad of Asp350-His471-Asp473;^43^ these catalytic residues, as well as numerous Ca^2+^-binding residues are retained between the human and *S. albulus* PADs (**Figure S6**). In contrast, the *P. gingivalis* (PAD_porph) is calcium-independent and cannot be aligned with the *S. albulus* PAD outside of short motifs near the active site (**Figure S7**). Moreover, few functional residues are conserved beyond those common to all members of the guanidino-modifying enzyme superfamily.^44,45^

The *S. albulus* PAD was used to generate a dataset of 837 bacterial PF03068 PADs using a PSI-BLAST query against all bacterial genomes in the NCBI non-redundant database (**Supplemental Dataset 1**).^46^ Protein accession identifiers were compiled, analyzed by RODEO,^21^ and then used to generate a sequence similarity network (SSN) and a maximum-likelihood phylogenetic tree to visualize the distribution of bacterial PADs belonging to PF03068 (**Figures 4** and **S8**).^47^ Homologs of the *S. albulus* PAD are present in diverse bacterial phyla, but are predominantly in Actinobacteria, Cyanobacteria, and Proteobacteria, with very few homologs in Bacteroidetes and Firmicutes. In contrast, members of PF04371 (PAD_porph), are more evenly distributed across prokaryotic phyla and are present in Actinobacteria, Proteobacteria, Firmicutes and Bacteroidetes, the latter including *P. gingivalis* (https://pfam.xfam.org/family/PF04371#tabview=tab7).^27^ Genomes that encode a PAD within these genera is fairly limited, as illustrated by only 1.6%, 0.1%, and 3.9% of sequenced actinobacteria, proteobacteria, and cyanobacteria, respectively, featuring an annotated homolog. Interestingly, the highest occurrence of annotated PADs is in Riflebacteria, which despite having only 11 genomes in the NCBI database, includes 15 annotated PADs. With a few notable examples, there is little evidence of horizontal gene transfer with PF03068, as proteins tend to cluster with other PADs from species within the same phylum (**Figures 4** and **S8**). Additionally, the %GC content of the genes encoding the PAD correlate strongly with the overall %GC content of the genome (**Figure S9**).^48^ These data indicate bacterial members of PF03068 are not disseminated across bacterial genomes due to recent horizontal gene transfer, although we cannot rule out the possibility that the genes were acquired by organisms having similar GC content. Notably, several proteobacterial PADs are more closely related to the actinobacterial PAD clade that includes those implicated in citrulassin Arg deimination. Intriguingly, most of these PAD-containing Proteobacteria are uncultured bacteria detected in aquatic and sediment metagenomes, raising questions concerning the genomic origins and ecology of PADs.

**Figure 3.**
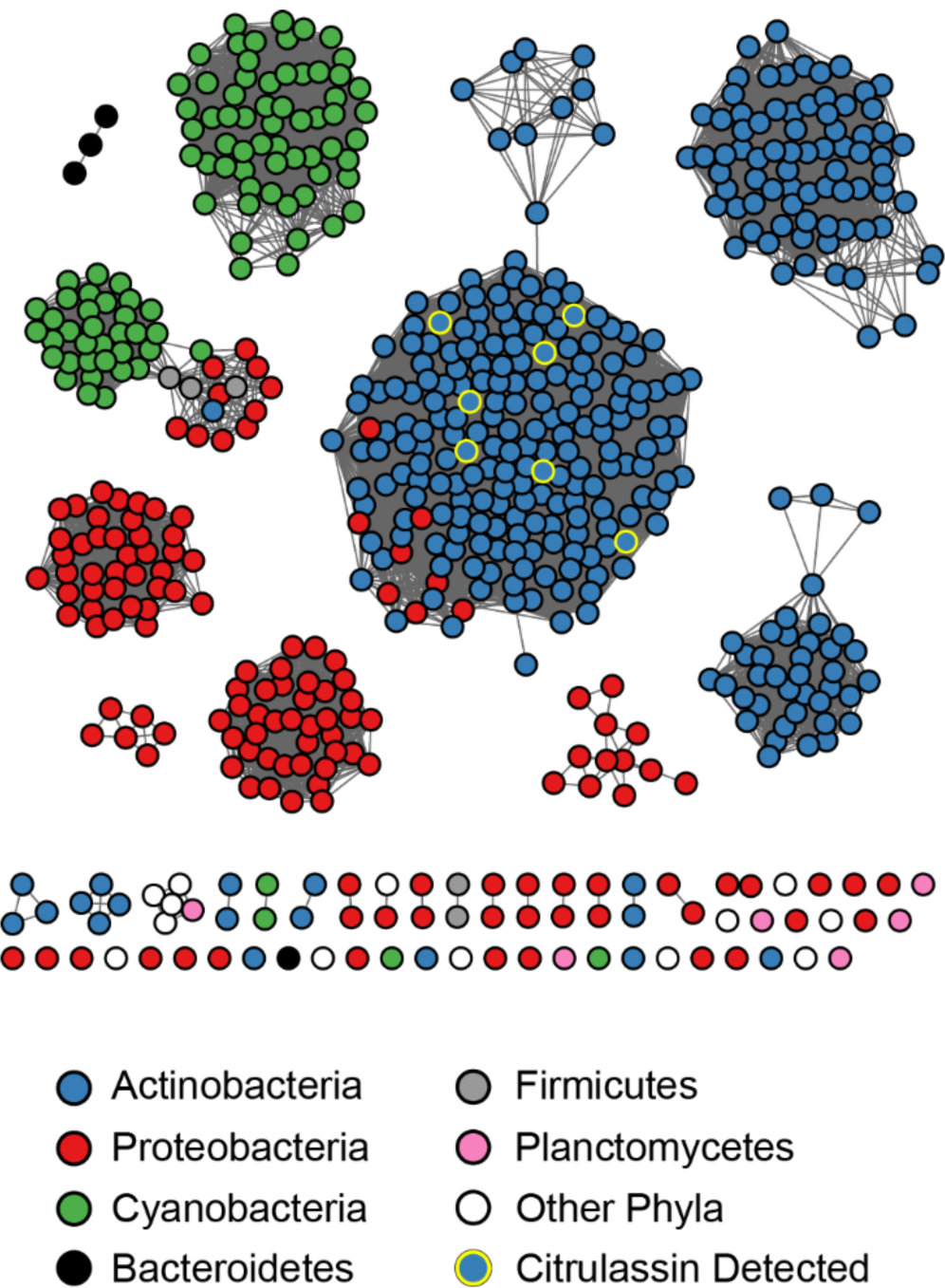
Protein sequence similarity network (SSN) of bacterial PADs. The SSN was generating using EFI-EST^47^, Sequences sharing 100% identity are conflated as a single node, and connected nodes indicate an alignment score of 125, see methods) and visualized by Cytoscape.^49^ Nodes are colored by phylum and detection of citrulassin (deiminated product) within the same strain.

**Figure 4:**
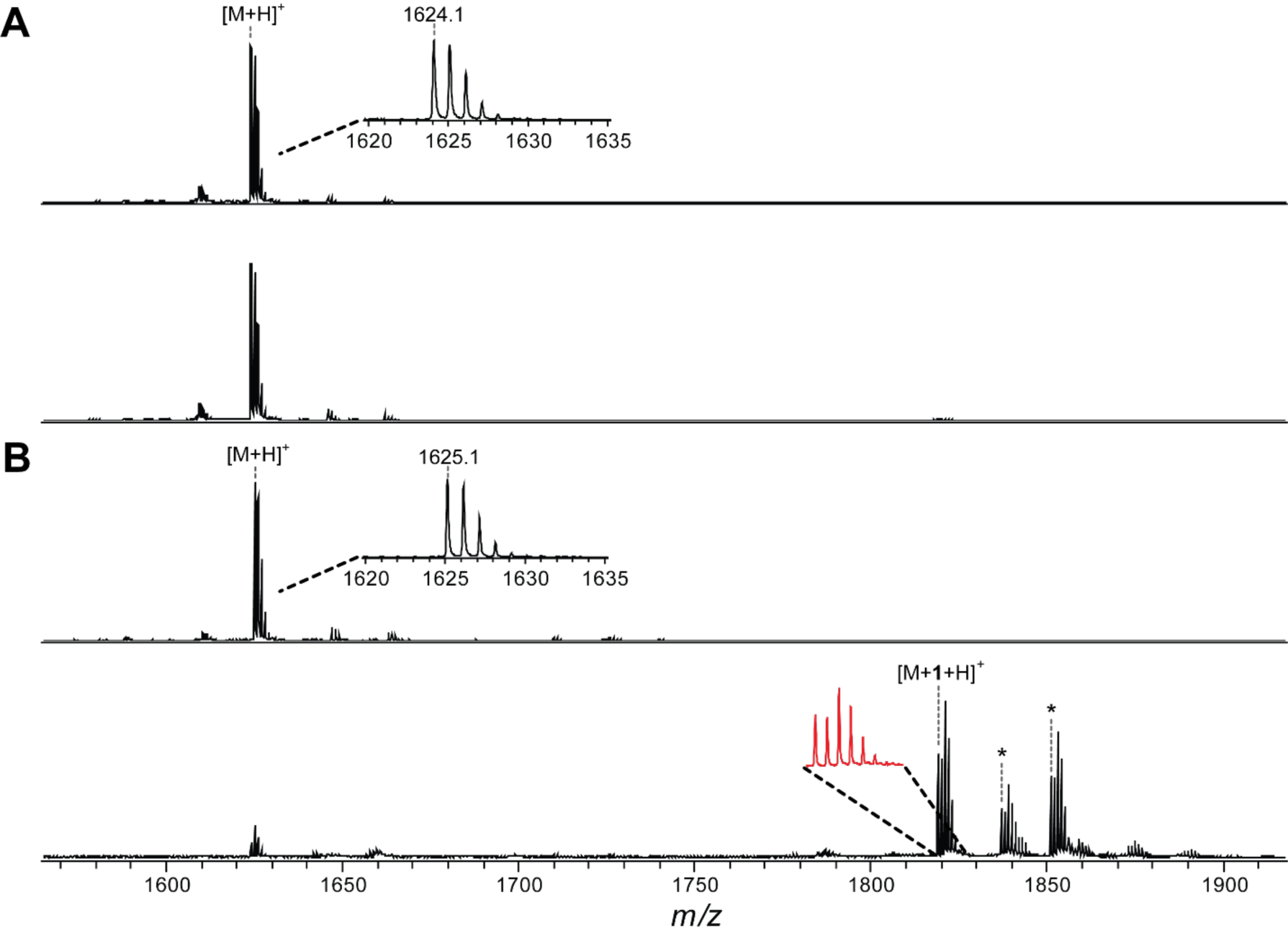
Citrulassin production in a *des*-citrulassin producer by PAD heterologous expression (**A**) MALDI-TOF mass spectrum of *Streptomyces katrae* extract containing *des*-citrulassin D unreacted (*top*) and reacted (*bottom*) with **1**. (**B**) MALDI-TOF mass spectrum of *S. katrae* extract expressing the PAD from *Streptomyces glaucescens*. Shown are samples unreacted (*top*) and reacted (*bottom*) with **1**. Insets show zoomed regions to visualize ^79^Br:^81^Br isotopic ratios. Asterisks denote solvated adducts corresponding to [M+ **1**+H2O+H]^+^ (left) and [M+**1**+MeOH+H]^+^ (right).

Co-occurrence analysis of citrulassin BGCs (± 8 coding sequences) shows little correlation beyond known lasso peptide biosynthetic proteins and furthermore demonstrates that PADs do not appear within lasso peptide biosynthetic operons (**Table S3**). Even PADs with a high degree of similarity appear in diverse genome neighborhoods, which do not bear resemblance to RiPP BGCs. The contexts within which the bacterial members of PF03068 appear are highly diverse (**Figure S10**). Thus, these PADs likely exert their activity on distally encoded substrate(s).

### PAD complementation in a *des*-citrulassin producer results in citrulassin production

The co-occurrence of PAD with the production of citrulassin provided strong, but ultimately circumstantial, support for the proposed role in Arg deimination. To further evaluate this putative biochemical role, a functional expression experiment designed to introduce the PAD-encoding gene from a verified citrulassin producer to the genome of a verified *des-*citrulassin producer that lacks a PAD-encoding gene. Using the *Streptomyces* phage φC31 integrase, plasmids containing an *attP* locus can be inserted into the chromosome of a recipient organism at the corresponding *attB* locus. Nearly all identified *des*-citrulassin producer genomes contain a φC31 *attB*, enabling *Escherichia coli*-*Streptomyces* conjugative transfer.^50,51^ The integrative plasmid places the PAD under strong constitutive promotion (*ermE*p*) to facilitate a higher level of expression within the heterologous host.^51, 52^ The *S. glaucescens* PAD was integrated into *Streptomyces katrae* NRRL B-16271, a native producer of *des*-citrulassin D (**Figure 4**, donor and recipient were arbitrarily chosen). Exconjugants were isolated by interative cultivation on a selective medium and chromosomal insertion of the PAD gene was confirmed via DNA sequencing (**Figure S11**). Successfully integrated strains were then grown on a medium that elicited *des*-citrulassin D production in wild-type *S. katrae*. Metabolites were extracted with MeOH and subsequently analyzed via MALDI-TOF-MS. PAD integration in *S. katrae* resulted in the production of a ion consistent with citrulline-containing citrulassin D (**Figure 4**). Reaction with **1** resulted in complete labeling, while wild-type *des*-citrulassin D was not labeled (**Figure 4**), suggesting that PAD integration had resulted in the conversion of the Arg within *des*-citrulassin D to citrulline. HRMS/MS confirmed that the produced lasso peptide was 0.98 Da heavier than expected for *des*-citrulassin D and elicited a fragmentation pattern consistent with the sequence of citrulassin D, featuring Arg9 converted to citrulline. Notably, the site-specific deimination of Arg9 of *des*-citrulassin D over Arg4 confirms our previously assigned site of deimination on citrulassin and demonstrates that expression of PAD is necessary and sufficient for conversion of Arg to citrulline in the context of citrulassin biosynthesis. Future work will be necessary to uncover the origins of the selectivity and the timing of deimination during citrulassin biosynthesis.

## Conclusion

Advances in genomic sequencing and bioinformatics have provided a foundation for a renaissance in NP discovery using genome-guided techniques, previously illustrated by the discovery of numerous new lasso peptides, including the citrulline-containing citrulassins. The versatility of genome-guided natural product discovery is strongly complemented by reactivity-based screening which exploits chemoselective targeting of organic functional groups to facilitate metabolite identification and isolation. In this work, we expand the toolkit of RBS to include primary ureido groups (i.e. citrulline) through their reactivity towards glyoxal-based probes, which, under mildly acidic conditions, are largely inert towards guanidino groups (i.e. Arg). Harnessing this reactivity led to the discovery of 11 new citrulassins and provided a means to perform comparative genomic analysis to discover the origins of the previously enigmatic Arg deimination in citrulassin biosynthesis. These analyses suggested that a family of bacterial PADs, divergent from the better-known bacterial PAD from *P. gingivalis*, were responsible for citrulassin deimination despite a lack of co-genomic context conservation. We confirmed the Arg deimination activity of this PAD converting a *des-*citrulassin producer into a citrulassin producer through heterologous expression of a PAD. This work highlights the synergy between genome- and reactivity-guided discovery approaches and provides a platform for identifying additional bacteria that produce NPs featuring the rare, non-proteinogenic primary ureido functional group. More broadly, we expect this workflow to be amenable towards uncovering both the prevalence and enzymatic origins of other reactive moieties present in RiPPs and other NPs.

## ASSOCIATED CONTENT

The Supporting Information is available free of charge via the web. This document contains a description of experimental methods, mass spectral characterization of newly discovered citrulassins, and bioinformatic analysis of the citrulassins and PAD enzymes.

## AUTHOR INFORMATION

### Funding sources

This work was supported by a grant from the National Institutes of Health (GM123998 to D.A.M.) and the Seemon Pines Fellowship from the Department of Chemistry at the University of Illinois at Urbana-Champaign (to G.A.H.). Funds to purchase the Bruker UltrafleXtreme MALDI TOF/TOF mass spectrometer were from the National Institutes of Health (S10 RR027109 A).

### Notes

The authors declare no competing financial interest.

## ACKNOWLEDGEMENT

We are grateful to W. W. Metcalf for access to many of the actinobacterial strains screened in this study, as well as the *E. coli* strains used for bacterial conjugation. P. Patel provided the script for automated % GC analysis.

## Notes

### Competing Interest Statement

The authors have declared no competing interest.

