## Supplementary material for "Reactivity-based screening for citrulline-containing natural products reveals a family of bacterial peptidyl arginine deiminases": SI document

[†]Current address: Leidos, National Energy Technology Laboratory, Pittsburgh, PA

**Methods**

**Materials.** All chemicals and solvents were purchased from Sigma-Aldrich, VWR, Ark Pharm, or Fisher Scientific and used without further purification unless otherwise noted.

**Growth and Extraction of Bacterial Cultures.** Actinomycete starter cultures (5 mL) were inoculated in liquid ATCC 172 growth medium (20 g/L soluble starch, 10 g/L glucose, 5 g/L yeast extract, 5 g/L N-Z amine, 1 g/L calcium carbonate) at 30 °C on a tube roller for 2-4 d. If necessary, glass beads were used to break apart mycelium aggregates before inoculation of plates. Approximately 500 µL of the seed culture was used to inoculate 10 cm agar plates (25 mL solid media per plate) of ISP2 (4 g/L yeast extract, 10 g/L malt extract, 4 g/L glucose, 15 g/L agar, pH 7.2), GUBC (6.25 g/L glycerol, 10 g/L sucrose, 10 g/L beef extract, 5 g/L casamino acids [Bacto], 20 mM phosphate buffer, and 10 mL of a solution of Balch's vitamins [described below], 15 g/L agar, pH 7.2), CFood (25 g/L glucose, 15 g/L fish meal [Dr. Earth], 2 g/L yeast extract, 4 g/L calcium carbonate, 15 g/L agar, pH 7.2), V8 (200 mL/L V8 vegetable juice, 3 g/L calcium carbonate, 15 g/L agar, pH 7.2), MS (20 g/L mannitol, 20 g/L soya flour [Kinako], 10 mM magnesium chloride, 15 g/L agar, pH 7.2) and ISP4 (10 g/L soluble starch, 1 g/L potassium phosphate dibasic, 1 g/L magnesium sulfate heptahydrate, 1 g/L sodium chloride, 2 g/L ammonium sulfate, 2 g/L calcium carbonate, 1 mg iron (II) sulfate heptahydrate, 1 mg zinc sulfate heptahydrate, 1 mg manganese (II) chloride, 15 g/L agar, pH 7.2) media. To the media specified was added Balch's vitamins[1], which consists of 2 mg biotin, 2 mg folic acid, 10 mg pyridoxine hydrochloride, 5 mg thiamine hydrochloride, 5 mg riboflavin, 5 mg nicotinic acid, and 5 mg DL-calcium pantothenate in 100 mL deionized water. Agar plates were grown at 30 °C for 10 d. Cell mass and the top layer of agar were scraped from the plate with a sterile razor blade and placed into a 1.5 mL Eppendorf tube. The cell mass was extracted with HPLC grade methanol for 2 h with occasional shaking. Solid material was removed by centrifugation (17,000 × g, 15-30 min) followed by removal of the liquid extract. Extracts were typically screened fresh (< 24 h), and without concentration. In cases where the signal from matrix-assisted laser desorption/ionization time-of-flight mass spectrometry (MALDI-TOF-MS) was poor, the extracts were concentrated under reduced pressure, reconstituted in water, and desalted using a C18 ZipTip (EMD Millipore) according to manufacturer specifications, and eluted into 80% aq. acetonitrile. Extracts were stored long term at -20 °C as a lyophilized powder.

**Phenylglyoxal Screening of Bacterial Extracts.** Under the optimized conditions, 10 µL of 10 mM bromophenylglyoxal hydrate (**1**) in 2% (*v/v*) methanolic trifluoroacetic acid was added to 10 µL bacterial extract in a 1.5 mL Eppendorf tube. Reactions were heated to 37 °C for 1 h, and then evaporated under reduced pressure using a speed vaccum concentrator (Thermo Scientific). Reactions were reconstituted in 10 µL HPLC grade methanol for further analysis. Reactions were subjected to MALDI-TOF-MS in reflector positive using alpha-cyano-4-hydroxycinnamic acid (CHCA) as a matrix. Spectra were analyzed manually for peaks displaying an isotope pattern consistent with one bromine. Initial hits were verified with a second reaction to assure reproducibility.

**High-Resolution Mass Spectrometry.** Lasso peptides of interest were either analyzed by high-resolution mass spectrometry (HRMS) as pure lyophilized powder, or desalted using a ZipTip according to manufacturer specifications, eluted using 80% aq. acetonitrile with 1% acetic acid, and subjected to centrifugation (17,000 × g, 10 min). Samples were infused onto a ThermoFisher Orbitrap Fusion Electrospray ionization mass spectrometer (ESI-MS) using an Advion TriVersa NanoMate. The MS was calibrated weekly using calibration mixture, following manufacturer instructions, and tuned daily with Pierce LTQ Velos ESI Positive Ion Calibration Solution (ThermoFisher). Spectra were collected in profile mode with a resolution of 100,000. Ions were selected for fragmentation in the Ultra-High-Field Orbitrap Mass Analyzer using an isolation width of 5 m/z, a normalized collision energy of 35, an activation q value of 0.4, and an activation time of 30 ms. Data analysis was performed using Thermo Xcalibur software.

**General Citrulassin Isolation.** In cases where desalting of small scale extracts provided insufficient material for HRMS, bacteria grown on 0.5 L of solid media were used to purify each citrulassin. The agar was cubed, frozen, thawed, and squeezed (filtering through cotton) to collect the bulk aqueous fraction. The squeezed agar was then extracted with 0.5 L of methanol for 2 h. The methanol was removed under vacuum and the remaining extract was combined with the liquid from the agar squeeze. The combined aqueous extract was vacuum filtered through Whatman filter paper to remove residual solid material. The extract was loaded onto a HyperSep C18 10 g SPE column (Thermo Scientific) pre-equilibrated with 100 mL acetonitrile, 100 mL 50% aq. acetonitrile, and 200 mL water before loading. After loading the unpurified extract, the column was washed with an additional 200 mL of water. Product was eluted with successive 50 mL portions of 10%, 20%, 30%, 40%, and 50% aq. acetonitrile, followed by 100 mL of acetonitrile. Fractions were analyzed via MALDI-TOF-MS, and fractions containing the citrulassin were pooled and dried under reduced pressure. The resulting crude material was redissolved in 80% aq. acetonitrile with 0.1% triflouroacetic acid, which was then injected onto a RediSep Rf SCX column (50 g medium, 60 Å pore size, 40–63 µm particle size, 230–400 mesh) and purified over a gradient of 0-100% 80% aq. acetonitrile with 0.1% triflouroacetic acid and 0.5 M potassium chloride (40 column volumes). Fractions were analyzed via MALDI-TOF-MS, pooled, and dried under reduced pressure. The partially purified material was then purified using a PerkinElmer Flexar HPLC equipped with a Betasil C18 (Thermo Scientific) reverse phase column (250 × 4.6 mm, 5 µm particle size, 100 Å pore size). A mobile phase of 10 mM aq. $NH_4HCO_3$/acetonitrile was used at a flow rate of 1 mL/min with a method of: 5% acetonitrile (isocratic, 5 min), 5-50% acetonitrile (gradient, 30 min), 50% acetonitrile (isocratic, 5 min) 50-100% acetonitrile (gradient, 5 min), 100% acetonitrile (isocratic, 5 min). Absorbance of the elution was monitored at 220 nm and fractions containing the citrulassin were dried under reduced pressure, and then used in HRMS.

**PAD and Citrulassin Bioinformatics.** The *Streptomyces albulus* NRRL B-3066 PAD (WP_064069847.1) was used as a query for Position-specific Iterated Basic Local Alignment Search Tool (PSI-BLAST)[2] limited to bacteria and an E-value of $10^{-10}$, returning 837 accession IDs. These sequences were submitted to the Enzyme Function Initiative Enzyme Similarity Tool (EFI-EST, https://efi.igb.illinois.edu/efi-est),[3] and sequences between 500 and 800 amino acids in length were used to generate a sequence similarity network (SSN) with an alignment score threshold of 150. Sequences that were 100% identical were conflated to a single node. The resulting SSN was visualized with Cytoscape (version 3.6.1)[4] and annotated with phyla information via Adobe Illustrator. The full list of PADs was submitted to Rapid ORF Description and Evaluation Online (RODEO; http://ripp.rodeo) for co-occurrence analysis and used to calculate the %GC of the PAD gene and organism genome. The PAD sequences were aligned using Multiple Alignment using Fast Fourier Transform (MAFFT, version 7.313) using the G-INS-i iterative refinement method.[5] The resulting alignment was used to create a maximum likelihood phylogenetic tree using FastTree (version 2.1.11)[6] and the tree visualized using the Interactive Tree of Life (iTOL) web tool (http://itol.embl.de/).[7] The citrulassin A lasso cyclase (WP_079136914.1) was used as a protein BLAST query with an E-value cutoff of $10^{-200}$. The resulting 191 NCBI accession IDs were submitted to RODEO for precursor identification and co-occurrence analysis. The citrulassin sequence logo was generated using WebLogo (http://weblogo.berkeley.edu/logo.cgi).[8]

**Spore Preparation of *Streptomyces katrae*.** *Streptomyces katrae* NRRL B-16271 was grown in 250 mL of ATCC 172 medium at 30 °C until high density (approximately 2-4 d). Cell mycelia were then plated onto 10 cm petri dishes of V8 medium and allowed to grow at 30 °C for 7 d. Spores were harvested with the addition of 5 mL sterile water with 0.1% (*v/v*) Tween 20 and gently scraped from the cell surface with a cell spreader. The liquid-spore suspension was filtered through sterile cotton using an autoclaved syringe and harvested by centrifugation (4,000 × g, 10 min, 4 °C). After centrifugation, the supernatant was discarded, and pellet resuspended in 2 mL sterile water with 25% (*v/v*) glycerol. This mixture was then flash frozen in liquid nitrogen and stored at -80 °C until use.

**Construction of Integrative Vector pAE4 for PAD Complementation in *Streptomyces katrae*.** Integrative vector pAE4 was modified by Gibson Assembly[9] to feature the strong constitutive promoter *ermE*p*.[10] Subsequent Gibson Assembly was used to insert the PAD from citrulassin E producer *Streptomyces glaucescens* downstream of the *ermE*p* promoter (NCBI accession identifier: WP_052413578.1). This vector was then transformed into conjugative *Escherichia coli* for insertion into *S. katrae* spores.

***E. coli-Streptomyces katrae* Conjugation.** *E. coli* conjugation strains WM6026 and WM6029 were transformed with pAE4-*ermE*p* constructs and selected on lysogeny broth (LB) plates containing 40 µg/mL 2,6-diaminopimelic acid (DAP) and 40 µg/mL apramycin sulfate (Apr). Colonies were selected for conjugation and inoculated in 5 mL of LB with 40 µg/mL DAP and 40 µg/mL Apr and allowed to grow at 37 °C for 16 h. Fresh LB with 40 µg/mL DAP and 40 µg/mL Apr was inoculated with 200 µL of the dense culture and incubated at 37 °C until OD600 = 0.6. Cells were then harvested by centrifugation (4,000 × g, 10 min, 4 °C) and the supernatant was discarded. The remaining cell pellet was resuspended and washed with 1 mL of 2×YT medium (16 g/L tryptone, 10 g/L yeast extract, 5 g/L NaCl). This resuspension and washing step was repeated three additional times, after which the pellet was resuspended in 1 mL of 2×YT medium with 40 µg/mL DAP and placed on ice until use for conjugation. As the *E. coli* conjugation strains reached OD600 = 0.6, a 500 µL frozen suspension of *S. katrae* spores was removed from -80 °C and heat activated at 56 °C for 15 min. Afterwards, 500 µL of 2×YT was added to the heat activated spore stock and mixed by pipetting. Conjugation between *E. coli* and *S. katrae* was initiated by aliquoting 200 µL of heat activated S. *katrae* spore mixture into three sterile 1.7 mL tubes along with 4 µL (50:1 *v/v*), 40 µL (5:1 *v/v*), and 200 µL (1:1 *v/v*) of *E. coli*. The spores and cells were mixed and then centrifuged (4,000 × g, 60 s, 25 °C) and incubated at 25 °C for 30 min. The cell pellet was then resuspended and plated on individual plates of V8 medium and incubated at 30 °C for 16 h. After 16 h, the plates were flooded with 2 mL of sterile-filtered 1 mg/mL aq. Apr solution and air-dried in a biosafety cabinet. The plates were then incubated at 30 °C until colonies appeared.

**Verifying Chromosomal Insertion of Conjugative Plasmid pAE4.** Individual colonies from 1 mg/mL Apr flooded V8 plates were inoculated onto plates of ISP2 medium with 40 µg/mL Apr. Individual colonies were picked from the ISP2 plates and grown in 5 mL ATCC 172 medium to high density. Genomic DNA was extracted using the Qiagen DNeasy UltraClean Microbial kit and checked for insertion of ermE*p-PAD by PCR and sequencing.

**Verifying Heterologous in vivo PAD Activity.** Individual colonies from 1 mg/mL Apr flooded V8 plates were restreaked on plates of ISP2 medium with 40 µg/mL Apr. Individual colonies were picked from the ISP2 plates and grown in 5 mL ATCC 172 medium to high density and were then inoculated onto plates of V8 Oats medium (20% *v/v* V8 tomato juice, 30 g/L steel cut oats) and grown at 30 °C for 7 d. Wild-type *S. katrae* was also grown and inoculated similarly as a positive control. Production of *des*- or citrulassins was confirmed by MALDI-TOF-MS, phenylglyoxal reactivity, and HRMS/MS.

**Table S1: Primers used in this study.** F, forward; R, reverse.

| Primer | Sequence (5’ → 3’) |
|---|---|
| pAE4_PAD_Gibson_Insert_F | CGGAGCAACGGAGGTACGGACATGCGCTCACGTACGCGCATACACCC |
| pAE4_PAD_Gibson_Insert_R | CCTGGGAAAACGTGAAGCCCCGGTCAGCCCCTGGGTGCGGCCCA |
| pAE4_PAD_Gibson_Vector_F | TGGGCCGCACCCAGGGGCTGACCGGGGCTTCACGTTTTCCCAGG |
| pAE4_PAD_Gibson_Vector_R | GGGTGTATGCGCGTACGTGAGCGCATGTCCGTACCTCCGTTGCTCCG |
| ermE-PAD insert F | GGCGGCAACCCTCAGC |
| ermE-PAD insert R | CGCTGAAGGTTCTTCGTGTCG |
| ermE_SeqF | ACGCGGTCGATCTTGACGG |

**Figure S1: Scheme for the workflow of reactivity-based screening (RBS) used in this study.** Genomes are mined for citrulassin biosynthetic gene clusters. Selected strains are then grown and the presence of peptidic citrulline is monitored by reacting crude cellular extracts with probe **1**. With a sufficient number of strains where the citrulline and Arg chemotypes are known, partial phylogenetic profiling is then used to populate a list of candidate gene(s) correlating with citrulassin deimination.

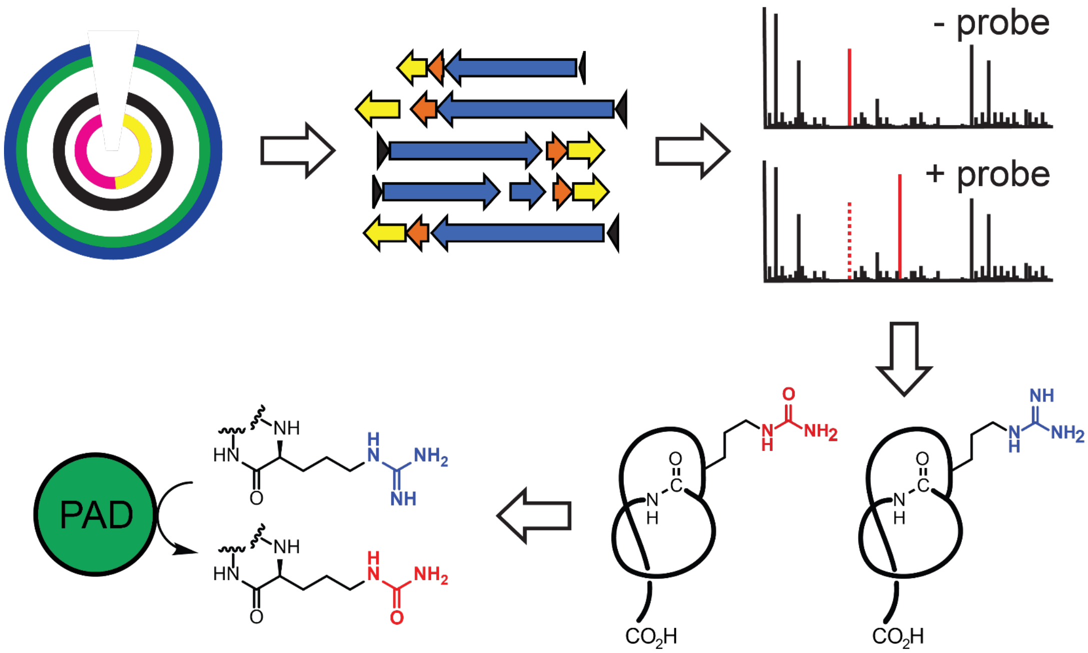

**Figure S2: 3-bromophenylglyoxal reacts with free citrulline but not arginine at low pH.** (A) Structure of citrulline. (B) Structure of arginine. (C) ESI mass spectra of free citrulline unreacted (*top*) or labeled with probe **1** (*bottom*). Labeled citrulline is magnified to show the characteristic $^{79}Br$:$^{81}Br$ isotope ratio. In the top spectrum, the ions marked with a * denotes $[Cit\text{-}OH+H]^+$, while those in the bottom spectrum are from left to right: $[probe\ hydrate\text{-}OH]^+$, $[Cit+probe\text{-}CO_2]^+$, and $[Cit+probe\text{-}OH+H]^+$ (D) MALDI-TOF mass spectra of free arginine prior to reaction with probe **1** (*top*) or after reaction with **1** (*bottom*, no observed reaction).

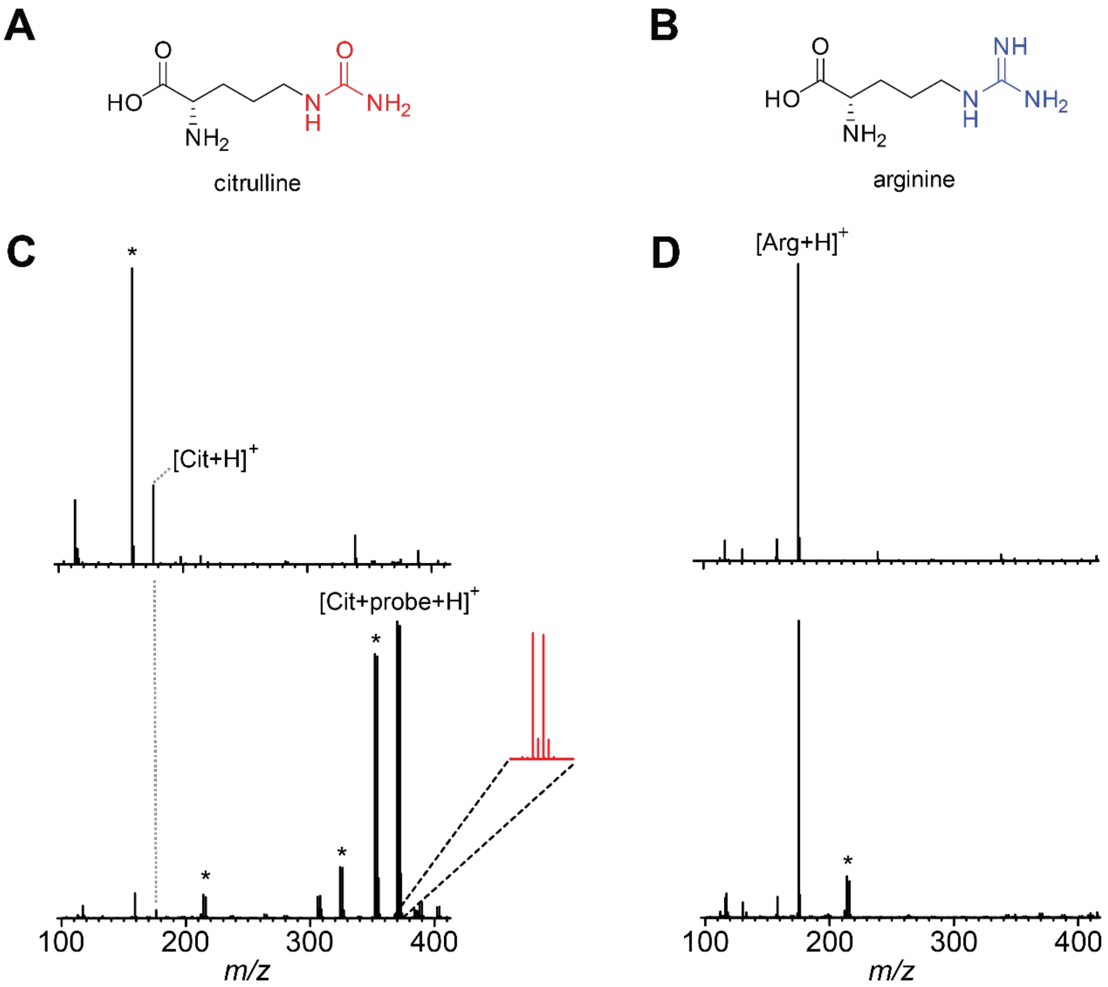

**Figure S3: 3-bromophenylglyoxal reacts with deimino-antipain but not antipain at low pH.** (A) Structure of *deimino*-antipain. (B) Structure of antipain. (C) MALDI-TOF mass spectra of *S. albulus* NRRL B-3066 extract unreacted (*top*) or labeled with probe **1** (*bottom*). The labeled *deimino*-antipain peak is magnified to show the characteristic $^{79}Br$:$^{81}Br$ isotope ratio. Ions marked with an asterisk are $[M+H_2O+H]^+$ (left) and $[M+MeOH+H]^+$ (right) (D) MALDI-TOF mass spectra of *S. lividans* 37C6 prior to reaction with probe **1** (*top*) and after reaction with **1** (*bottom,* no observed reaction).

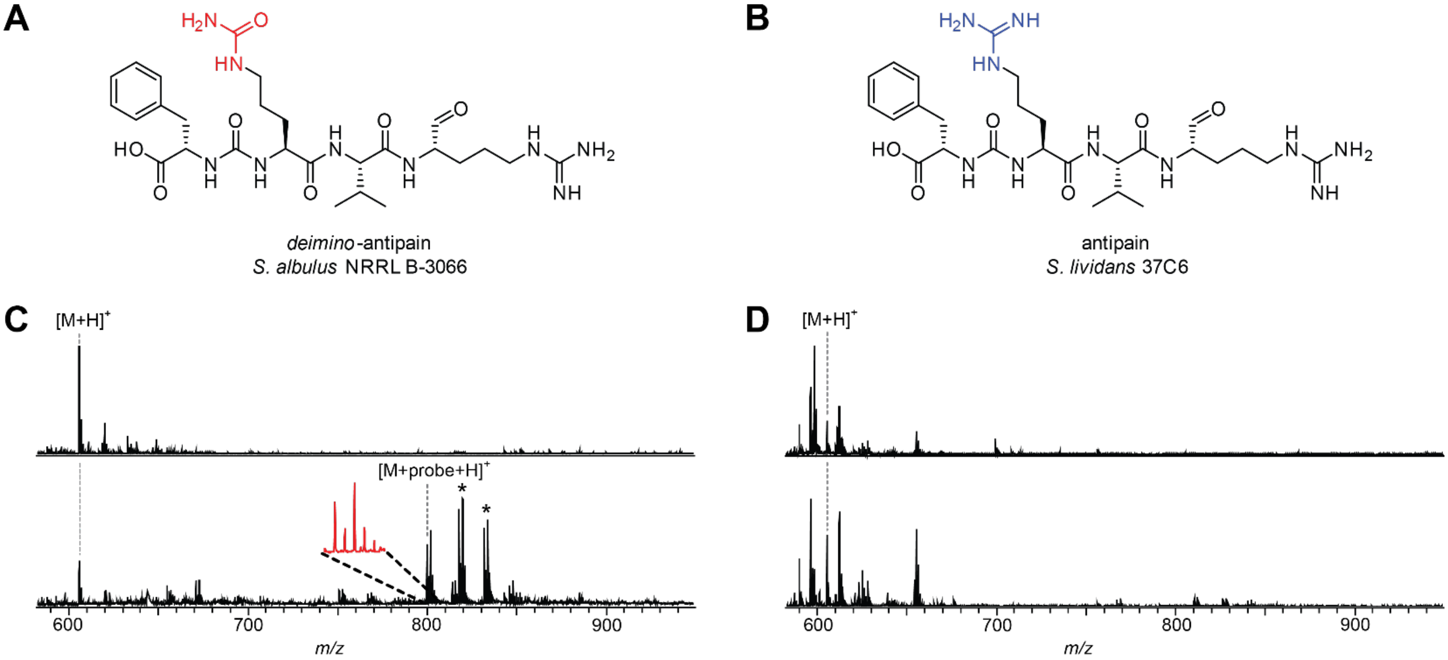

**Table S2: Citrulassins local co-occurrence analysis.** Co-occurrence analysis of ORFs ±8 from the citrulassin BGCs. Protein families (Pfams) with ≥10% co-occurrence are listed. Proteins involved in lasso peptide biosynthesis are bolded. A comprehensive list can be found in Supplementary Dataset 1. This analysis shows no evidence of lasso peptide biosynthetic gene co-occurrence with any gene encoding a PAD (PF03068).

| PfamID | Count | Percent | Name | Description |
|---|---|---|---|---|
| PF00005 | 192 | 106 | ABC_tran | ABC transporter |
| **PF00733** | **181** | **100** | **Asn_synthase** | **Asparagine synthase** |
| **PF05402** | **180** | **99** | **PqqD** | **Coenzyme PQQ synthesis protein D (PqqD)** |
| **PF13471** | **177** | **98** | **Transglut_core3** | **Transglutaminase-like superfamily** |
| PF02463 | 171 | 94 | SMC_N | RecF/RecN/SMC N terminal domain |
| PF00664 | 167 | 92 | ABC_membrane | ABC transporter transmembrane region |
| PF00440 | 42 | 23 | TetR_N | Bacterial regulatory proteins, tetR family |
| PF00196 | 33 | 18 | GerE | Bacterial regulatory proteins, luxR family |
| PF13304 | 32 | 18 | AAA_21 | AAA domain, putative AbiEii toxin, Type IV TA system |
| PF00795 | 29 | 16 | CN_hydrolase | Carbon-nitrogen hydrolase |
| PF13560 | 29 | 16 | HTH_31 | Helix-turn-helix domain |
| PF00561 | 27 | 15 | Abhydrolase_1 | alpha/beta hydrolase fold |
| PF07690 | 27 | 15 | MFS_1 | Major Facilitator Superfamily |
| PF00528 | 25 | 14 | BPD_transp_1 | Binding-protein-dependent transport system inner membrane component |
| PF01381 | 25 | 14 | HTH_3 | Helix-turn-helix |
| PF08281 | 25 | 14 | Sigma70_r4_2 | Sigma-70, region 4 |
| PF00106 | 23 | 13 | adh_short | short chain dehydrogenase |
| PF13561 | 23 | 13 | adh_short_C2 | Enoyl-(Acyl carrier protein) reductase |
| PF00583 | 22 | 12 | Acetyltransf_1 | Acetyltransferase (GNAT) family |
| PF12146 | 22 | 12 | Hydrolase_4 | Serine aminopeptidase, S33 |
| PF12697 | 22 | 12 | Abhydrolase_6 | Alpha/beta hydrolase family |
| PF13556 | 21 | 12 | HTH_30 | PucR C-terminal helix-turn-helix domain |
| PF07228 | 20 | 11 | SpoIIE | Stage II sporulation protein E (SpoIIE) |
| PF12802 | 20 | 11 | MarR_2 | MarR family |
| PF13384 | 18 | 10 | HTH_23 | Homeodomain-like domain |

**Figure S5: (parts i-xii) Mass spectrometry characterization of novel citrulassins.** (A) MALDI-TOF-MS spectra of producing organism methanolic extract unreacted (*top*) or labeled with probe **1** (*bottom*). (B) HRMS of either the $[M+H]^+$ or $[M+2H]^{2+}$ (indicated) was used to calculate exact mass and error from the expected molecular formula. (C) Collision-induced dissociation spectrum of the $[M+H]^+$ or $[M+2H]^{2+}$ ion. Labeled peaks correspond to identified b- and y-ions. (D) Schematized lasso peptide diagram with identified fragment ions (shown unthreaded to aid interpretation). (E) Table of masses of the observed b-and y-ions shown in panels B and C. Cit, citrulline; Orn, ornithine.

**Part i: *des*-citrulassin C, *Streptomyces tricolor* NRRL B-16925, ISP4 media**

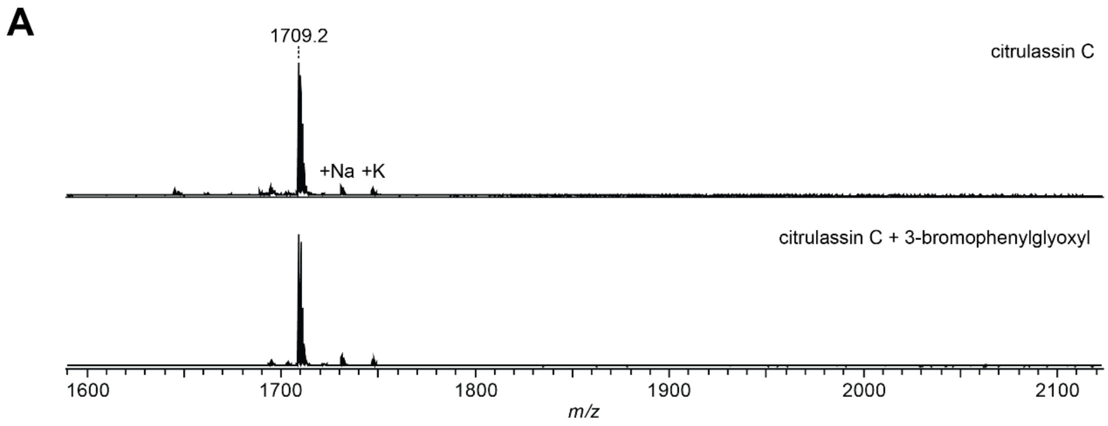

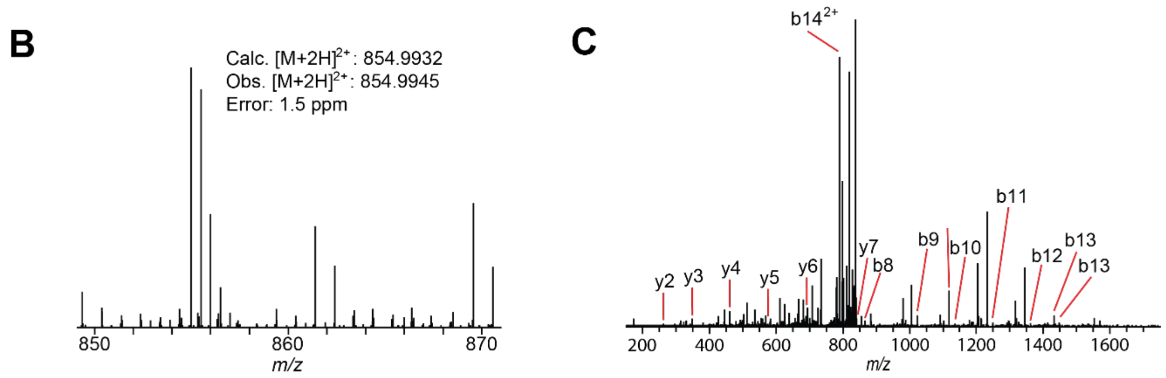

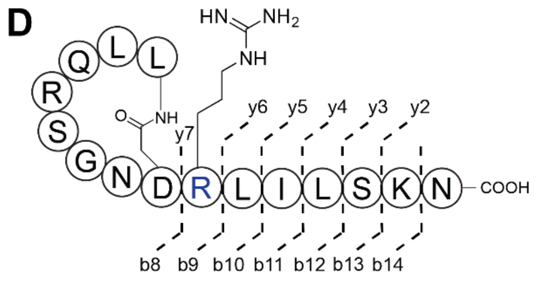

**E**

| Ion | Calc. Mass | Obs. Mass | Error (ppm) |
|---|---|---|---|
| **b14$^{2+}$** | 788.9677 | 788.9654 | 2.9 |
| **b13** | 1448.83322 | 1448.8279 | 3.7 |
| **b12** | 1361.8012 | 1361.7955 | 4.2 |
| **b11** | 1248.7171 | 1248.7133 | 3.0 |
| **b10** | 1135.6331 | 1135.6305 | 2.3 |
| **b9** | 1022.5490 | 1022.5460 | 2.9 |
| **b8** | 866.4479 | 866.4451 | 3.2 |
| **y2** | 261.1557 | 261.1552 | 1.9 |
| **y3** | 348.1878 | 348.1870 | 2.3 |
| **y4** | 461.2718 | 461.2707 | 2.4 |
| **y5** | 574.3559 | 574.3543 | 2.8 |
| **y6** | 687.4400 | 687.4380 | 2.9 |
| **y7** | 843.5411 | 843.5389 | 2.6 |

**Part ii: *des*-citrulassin D, *Streptomyces katrae* NRRL B-16271, V8 media**

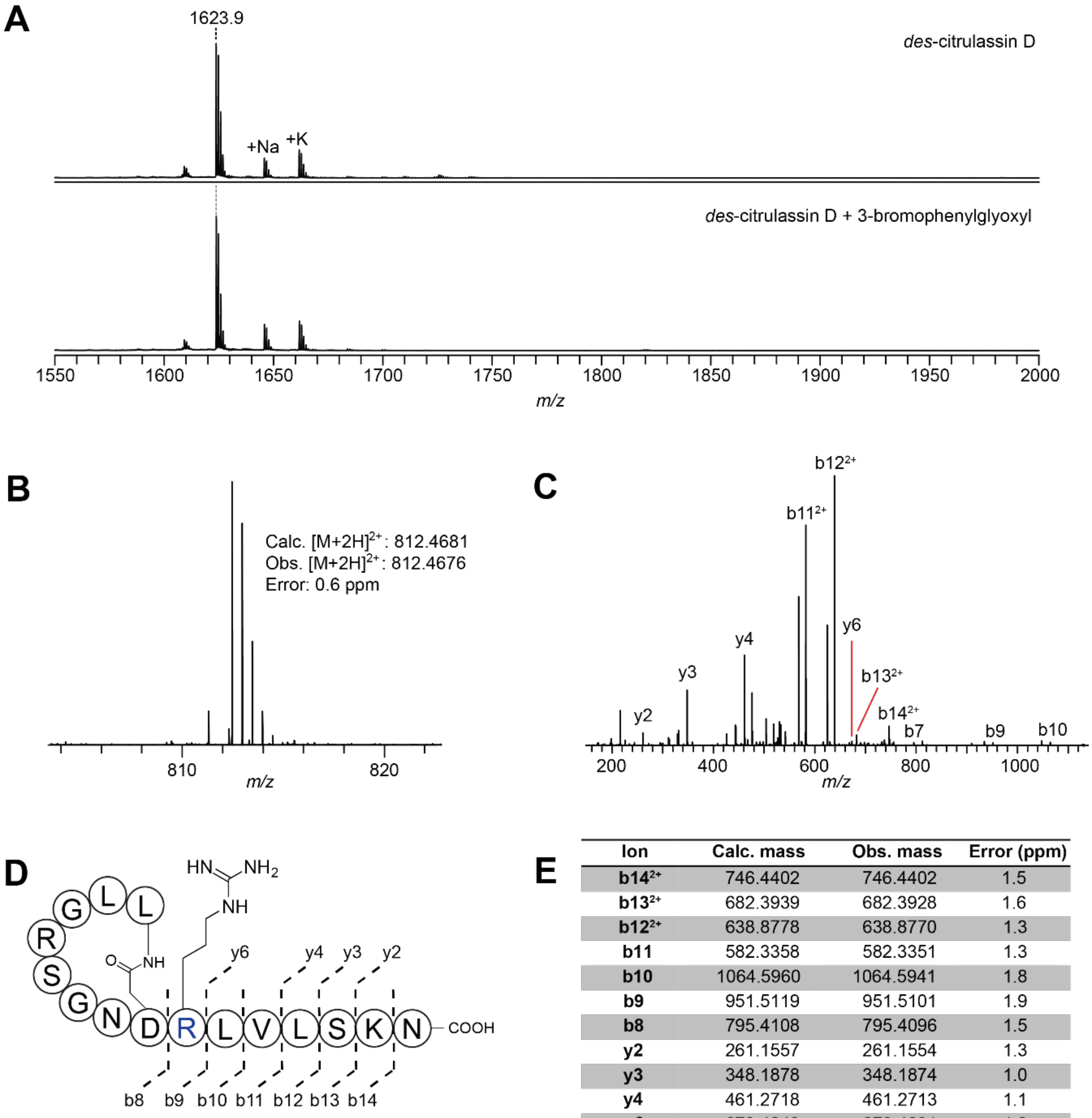

**E**

| Ion | Calc. mass | Obs. mass | Error (ppm) |
|---|---|---|---|
| **b14$^{2+}$** | 746.4402 | 746.4402 | 1.5 |
| **b13$^{2+}$** | 682.3939 | 682.3928 | 1.6 |
| **b12$^{2+}$** | 638.8778 | 638.8770 | 1.3 |
| **b11** | 582.3358 | 582.3351 | 1.3 |
| **b10** | 1064.5960 | 1064.5941 | 1.8 |
| **b9** | 951.5119 | 951.5101 | 1.9 |
| **b8** | 795.4108 | 795.4096 | 1.5 |
| **y2** | 261.1557 | 261.1554 | 1.3 |
| **y3** | 348.1878 | 348.1874 | 1.0 |
| **y4** | 461.2718 | 461.2713 | 1.1 |
| **y6** | 673.4243 | 673.4231 | 1.8 |

**Part iii: citrulassin D, *Streptomyces katrae* NRRL B-16271 PAD exconjugant, V8 media**

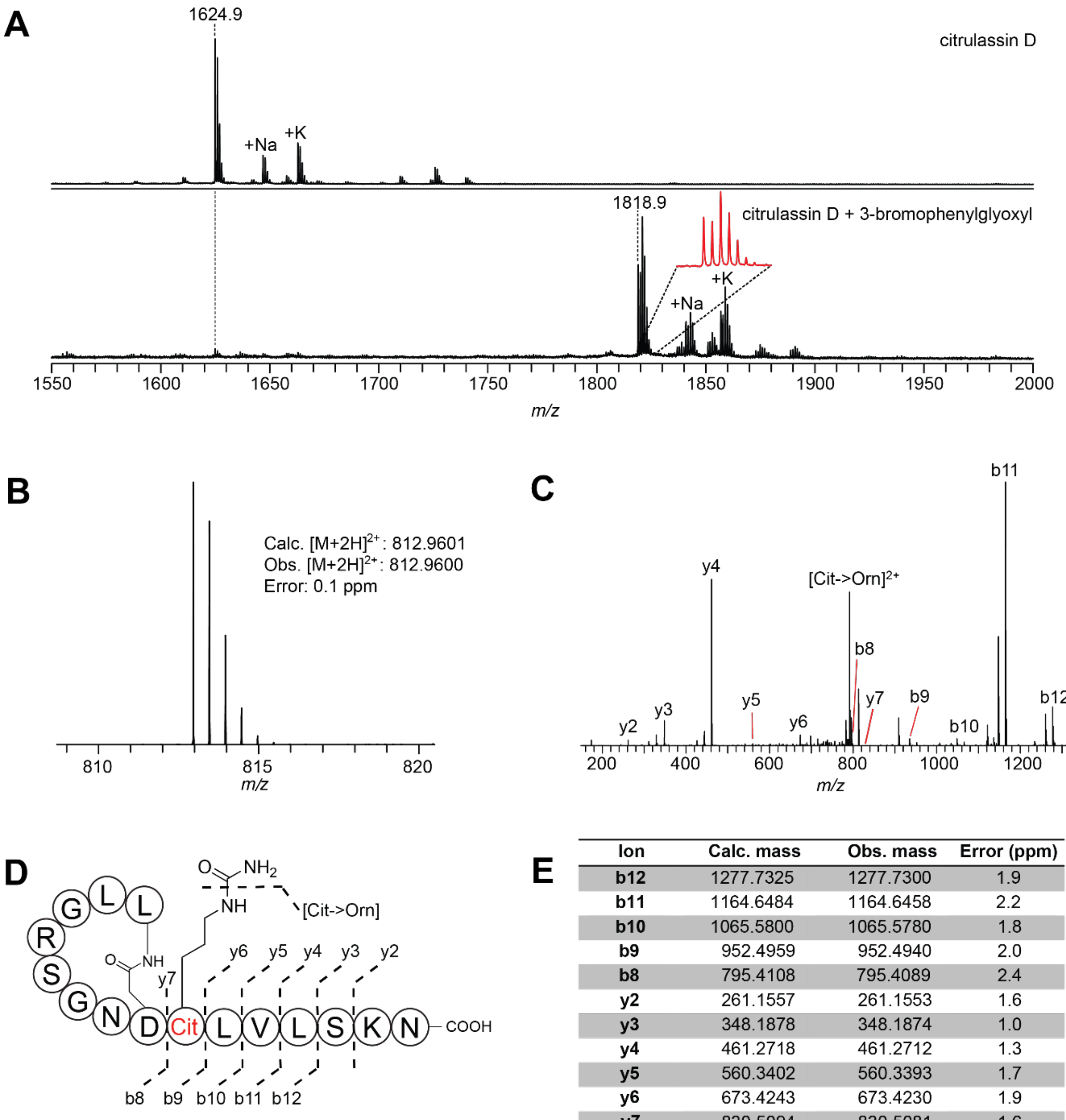

**E**

| Ion | Calc. mass | Obs. mass | Error (ppm) |
|---|---|---|---|
| **b12** | 1277.7325 | 1277.7300 | 1.9 |
| **b11** | 1164.6484 | 1164.6458 | 2.2 |
| **b10** | 1065.5800 | 1065.5780 | 1.8 |
| **b9** | 952.4959 | 952.4940 | 2.0 |
| **b8** | 795.4108 | 795.4089 | 2.4 |
| **y2** | 261.1557 | 261.1553 | 1.6 |
| **y3** | 348.1878 | 348.1874 | 1.0 |
| **y4** | 461.2718 | 461.2712 | 1.3 |
| **y5** | 560.3402 | 560.3393 | 1.7 |
| **y6** | 673.4243 | 673.4230 | 1.9 |
| **y7** | 830.5094 | 830.5081 | 1.6 |
| **[Cit-> Orn]$^{2+}$** | 791.4572 | 791.4561 | 1.4 |

**Part iv: citrulassin E, *Streptomyces glaucescens* NRRL B-11408, alternate MS media**

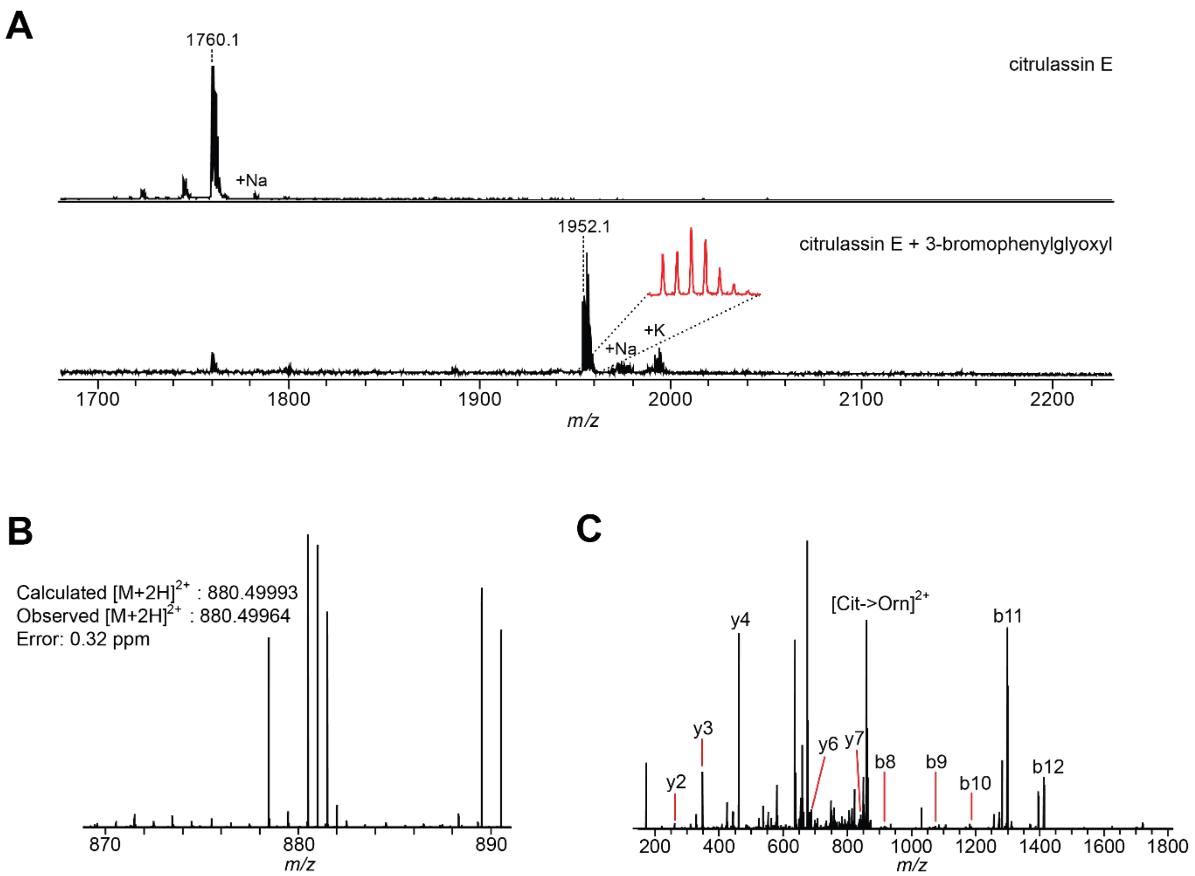

D

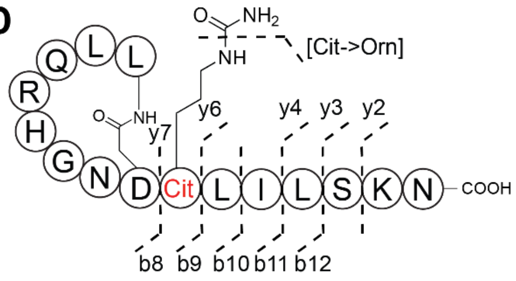

E

| Species | Calc. mass | Obs. Mass | Error (ppm) |
|---|---|---|---|
| **b12** | 1412.8121 | 1412.8087 | 2.4 |
| **b11** | 1299.7280 | 1299.7248 | 2.5 |
| **b10** | 1186.6440 | 1186.6413 | 2.2 |
| **b9** | 1073.5599 | 1073.5576 | 2.1 |
| **b8** | 916.4748 | 916.4732 | 1.7 |
| **y2** | 261.1557 | 261.1554 | 1.4 |
| **y3** | 348.1878 | 348.1871 | 1.8 |
| **y4** | 461.2718 | 461.2711 | 1.5 |
| **y6** | 687.4350 | 687.4388 | 1.7 |
| **y7** | 844.5251 | 844.5233 | 2.1 |
| **[Cit-> Orn]$^{2+}$** | 858.9970 | 858.9954 | 1.1 |

**Part v: *des*-citrulassin F, *Streptomyces avermitilis* NRRL B-16169, ISP4 media**

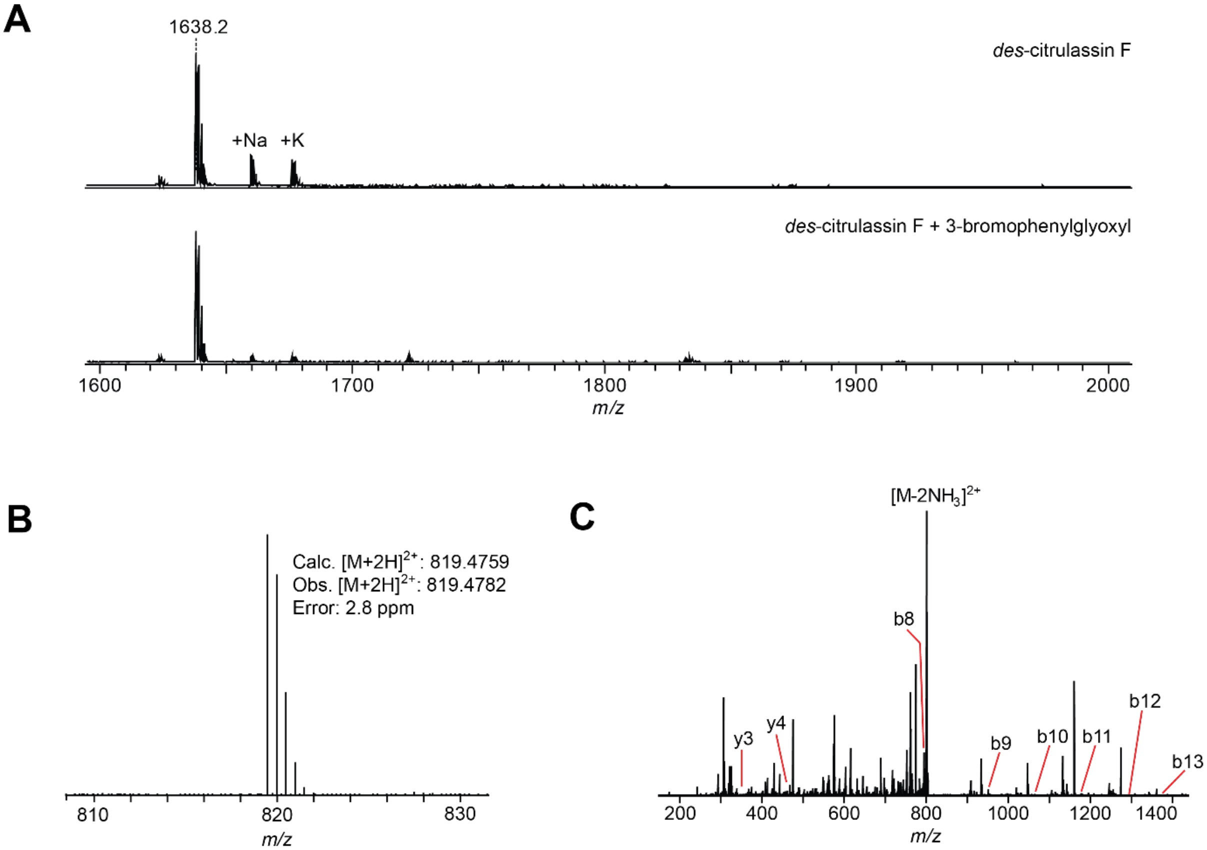

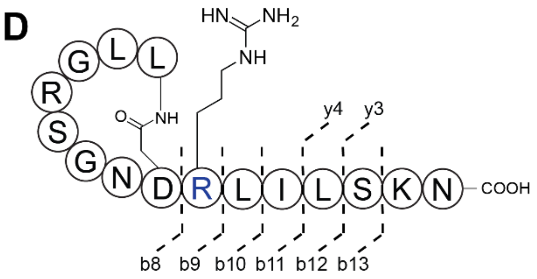

**E**

| Ion | Calc. mass | Obs. mass | Error (ppm) |
|---|---|---|---|
| **[M-2NH$_3$]$^{2+}$** | 801.4416 | 801.4415 | 0.1 |
| **b13** | 1377.7961 | 1377.7958 | 0.2 |
| **b12** | 1290.7641 | 1290.7656 | 1.2 |
| **b11** | 1177.6800 | 1177.6796 | 0.3 |
| **b10** | 1064.5960 | 1064.5949 | 1.0 |
| **b9** | 951.5119 | 951.5125 | 0.6 |
| **b8** | 795.4125 | 795.4108 | 2.1 |
| **y3** | 348.1878 | 348.1880 | 0.6 |
| **y4** | 461.2718 | 461.2721 | 0.7 |

**Part vi: citrulassin F, *Streptomyces torulosus* NRRL S-189, ISP4 media**

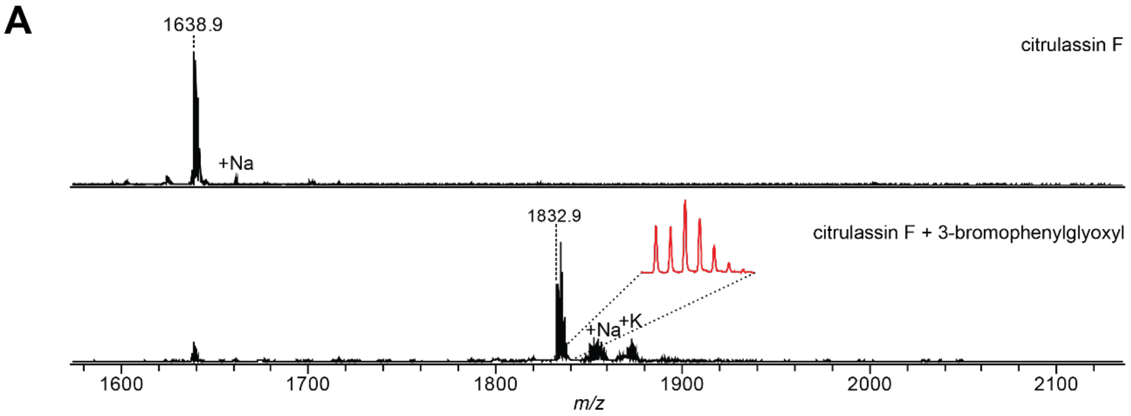

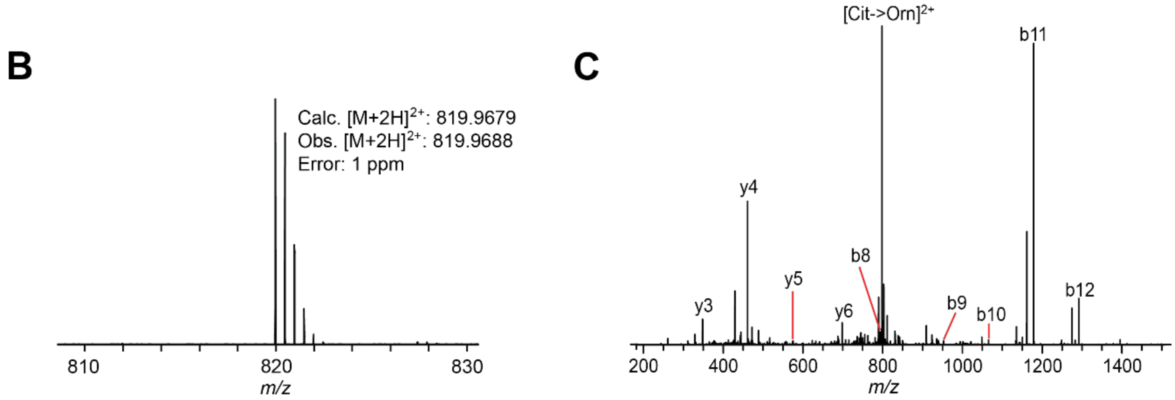

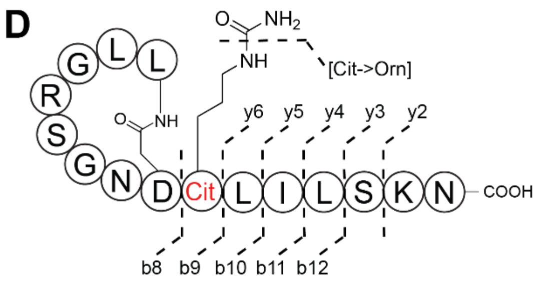

**E**

| Ion | Calc. mass | Obs. mass | Error (ppm) |
|---|---|---|---|
| b12 | 1291.7486 | 1291.7486 | 0 |
| b11 | 1178.6646 | 1178.6646 | 0 |
| b10 | 1065.5805 | 1065.5802 | 0.3 |
| b9 | 952.4965 | 952.4965 | 0 |
| b8 | 795.4113 | 795.4108 | 0.6 |
| y2 | 261.1557 | 261.1559 | 0.8 |
| y3 | 348.1878 | 348.1879 | 0.3 |
| y4 | 574.3559 | 574.3560 | 0.2 |
| y5 | 461.2718 | 461.2719 | 0.2 |
| y6 | 687.4400 | 687.4403 | 0.4 |
| [Cit-> Orn]$^{2+}$ | 798.4650 | 798.4660 | 1.3 |

**Part vii: *des*-citrulassin G, *Streptomyces auratus* NRRL 8097, ISP4 media**

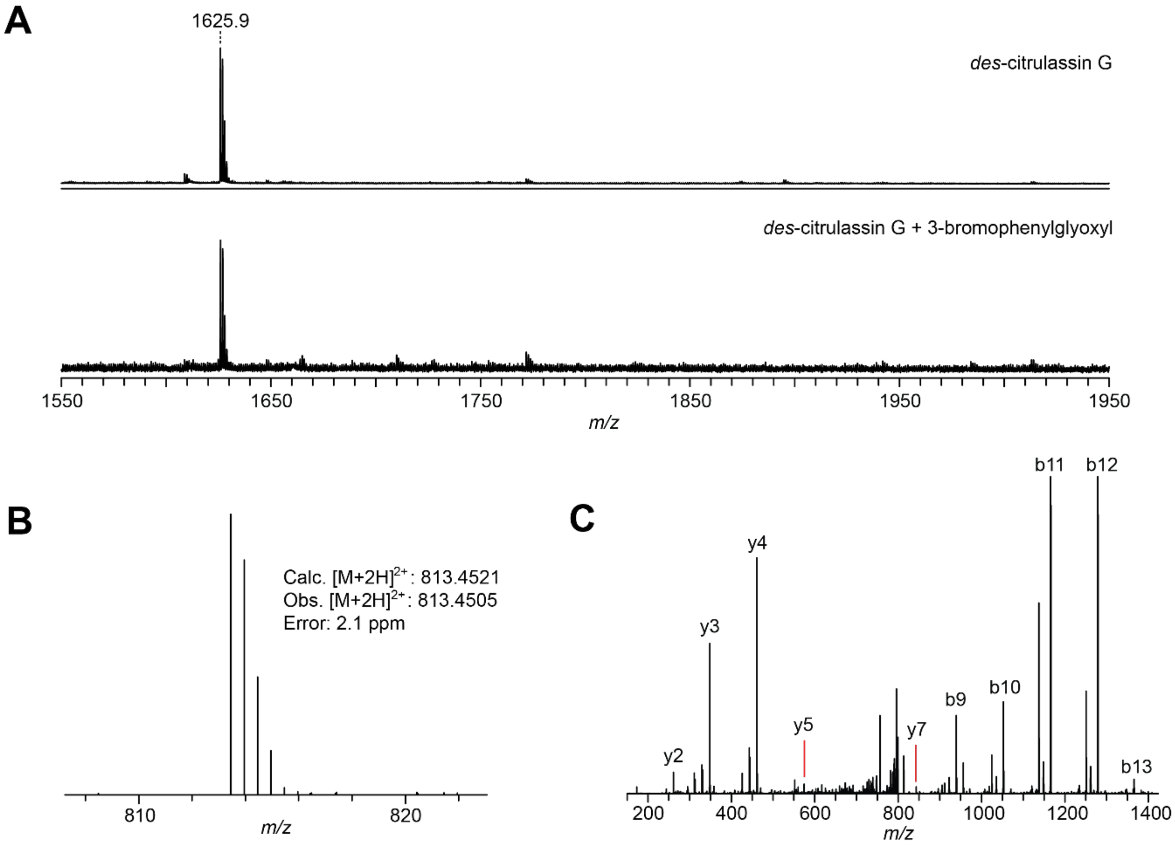

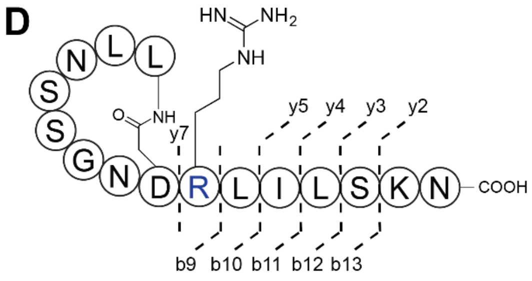

**E**

| Ion | Calc. mass | Obs. mass | Error (ppm) |
|---|---|---|---|
| **b13** | 1365.7485 | 1365.7453 | 2.3 |
| **b12** | 1278.7165 | 1278.7132 | 2.5 |
| **b11** | 1165.6324 | 1165.6295 | 2.5 |
| **b10** | 1052.5483 | 1052.5456 | 2.6 |
| **b9** | 939.4643 | 939.4617 | 2.8 |
| **y2** | 261.1557 | 261.1552 | 1.9 |
| **y3** | 348.1878 | 348.1871 | 1.9 |
| **y4** | 461.2718 | 461.2709 | 2.1 |
| **y5** | 574.3559 | 574.3546 | 2.3 |
| **y7** | 843.5411 | 843.5388 | 2.7 |

**Part viii: citrulassin H, *Streptomyces sp.* NRRL S-118, alternate MS media**

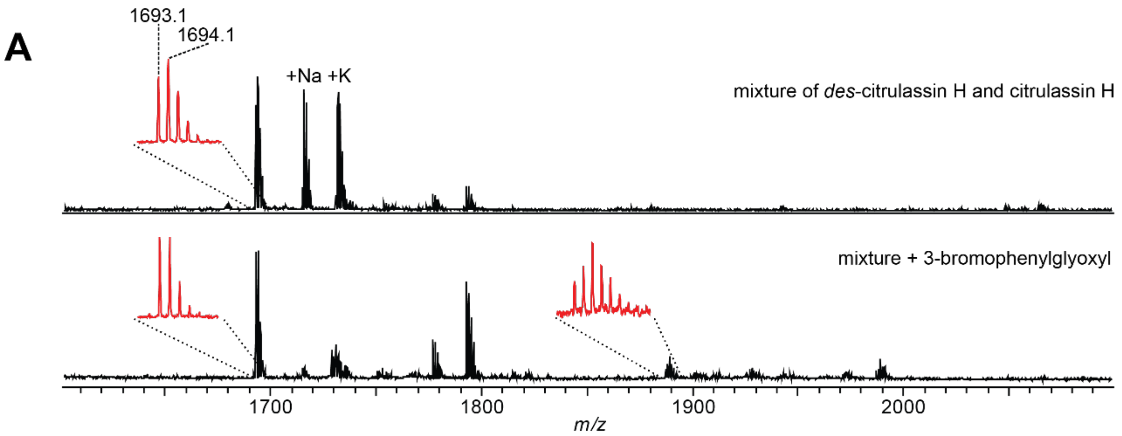

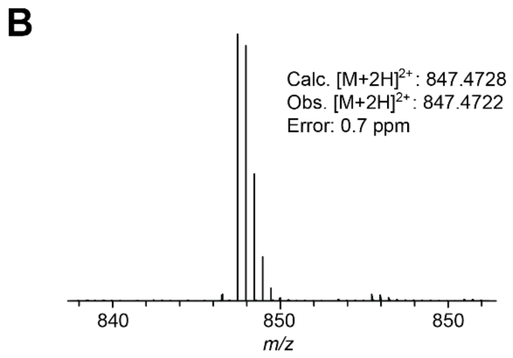

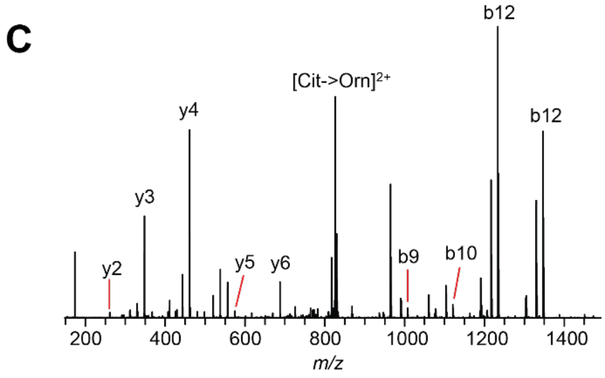

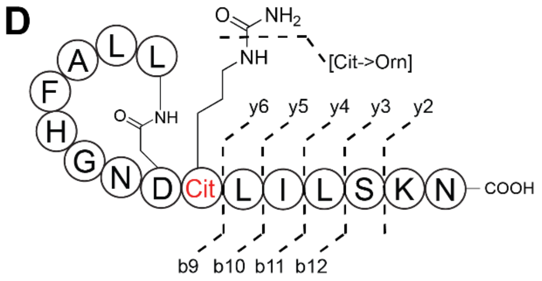

**E**

| Ion | Calc. mass | Obs. mass | Error (ppm) |
|---|---|---|---|
| **b12** | 1346.7579 | 1346.7544 | 2.6 |
| **b11** | 1233.6739 | 1233.6711 | 2.3 |
| **b10** | 1120.5898 | 1120.5873 | 2.2 |
| **b9** | 1007.5057 | 1007.5034 | 2.3 |
| **y2** | 261.1557 | 261.1552 | 2.0 |
| **y3** | 348.1878 | 348.1870 | 2.2 |
| **y4** | 461.2718 | 461.2709 | 2.0 |
| **y5** | 574.3559 | 574.3548 | 1.9 |
| **y6** | 687.4400 | 687.4386 | 2.0 |
| **[Cit-> Orn]$^{2+}$** | 825.9699 | 825.9680 | 2.3 |

**Part ix: *des*-citrulassin J, *Streptomyces natalensis* NRRL B-5314, alternate MS media**

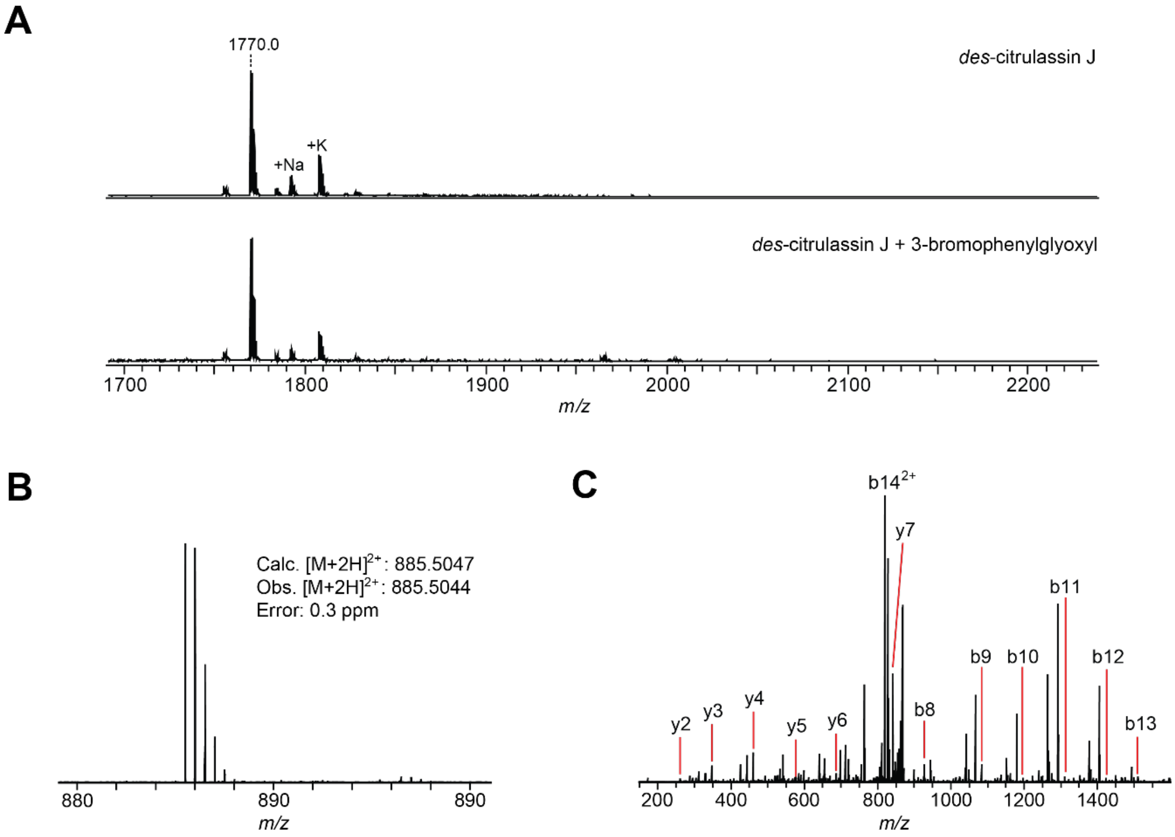

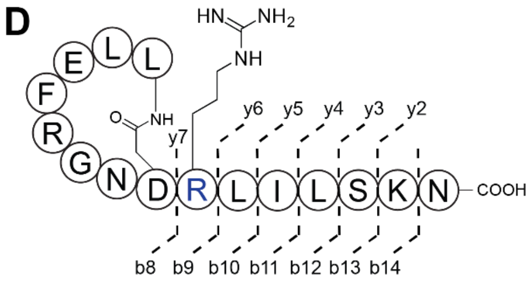

**E**

| Ion | Calc. mass | Obs. mass | Error (ppm) |
|---|---|---|---|
| **b14$^{2+}$** | 819.4779 | 819.4770 | 1.1 |
| **b13** | 1509.8536 | 1509.8519 | 1.1 |
| **b12** | 1422.8216 | 1422.8189 | 1.9 |
| **b11** | 1309.7370 | 1309.7355 | 1.1 |
| **b10** | 1196.6535 | 1196.6525 | 0.8 |
| **b9** | 1083.5694 | 1083.5684 | 0.9 |
| **b8** | 927.4683 | 927.4667 | 1.7 |
| **y2** | 261.1557 | 261.1557 | 0.0 |
| **y3** | 348.1878 | 348.1876 | 0.6 |
| **y4** | 461.2718 | 461.2714 | 0.9 |
| **y5** | 574.3559 | 574.3553 | 1.0 |
| **y6** | 687.4400 | 687.4393 | 1.0 |
| **y7** | 843.5411 | 843.5396 | 1.8 |

**Part x: *des*-citrulassin K, *Streptomyces sp.* S-920, V8 media**

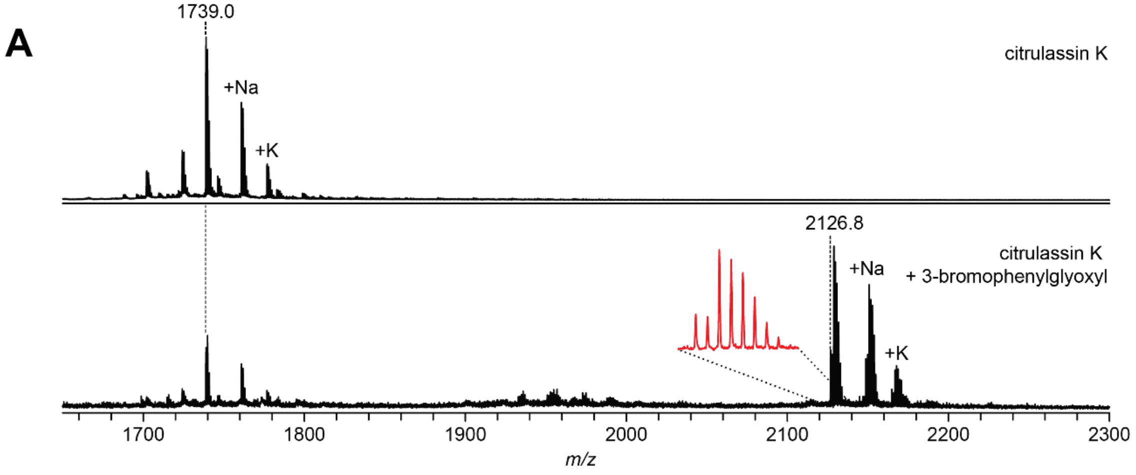

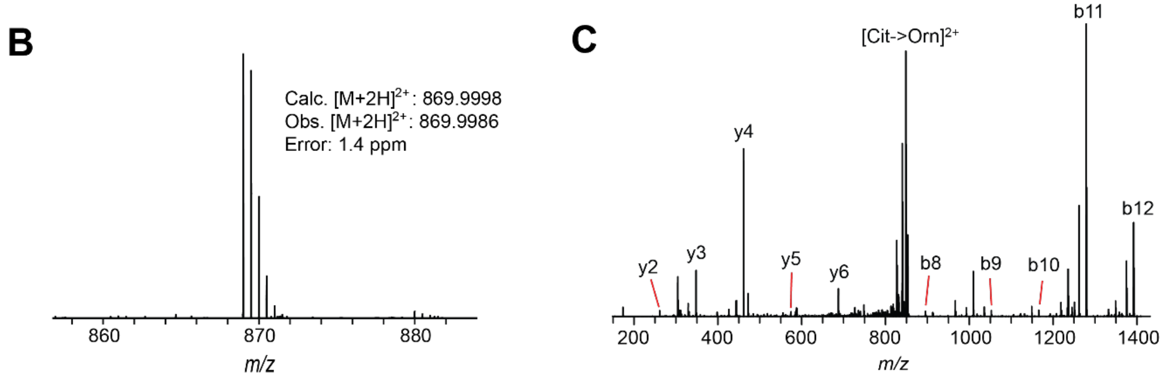

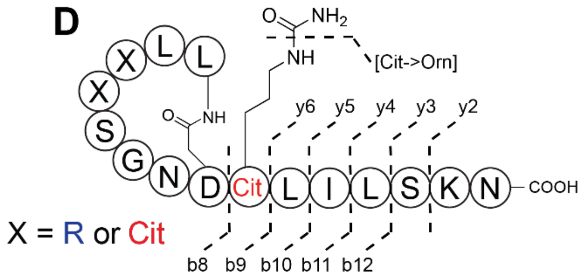

**E**

| Ion | Calc. mass | Obs. mass | Error (ppm) |
|---|---|---|---|
| **b12** | 1391.8118 | 1391.8078 | 2.9 |
| **b11** | 1278.7277 | 1278.7243 | 2.7 |
| **b10** | 1165.6436 | 1165.6408 | 2.4 |
| **b9** | 1052.5596 | 1052.5571 | 2.4 |
| **b8** | 895.4744 | 895.4724 | 2.2 |
| **y2** | 261.1557 | 261.1553 | 1.6 |
| **y3** | 348.1878 | 348.1871 | 1.9 |
| **y4** | 461.2718 | 461.2711 | 1.6 |
| **y5** | 574.3559 | 574.3547 | 2.1 |
| **y6** | 687.4400 | 687.4385 | 2.1 |
| **[Cit-> Orn]$^{2+}$** | 848.4969 | 848.4948 | 2.5 |

**Part xi: *des*-citrulassin L, *Streptomyces sp.* S-481, alternate MS media**

A

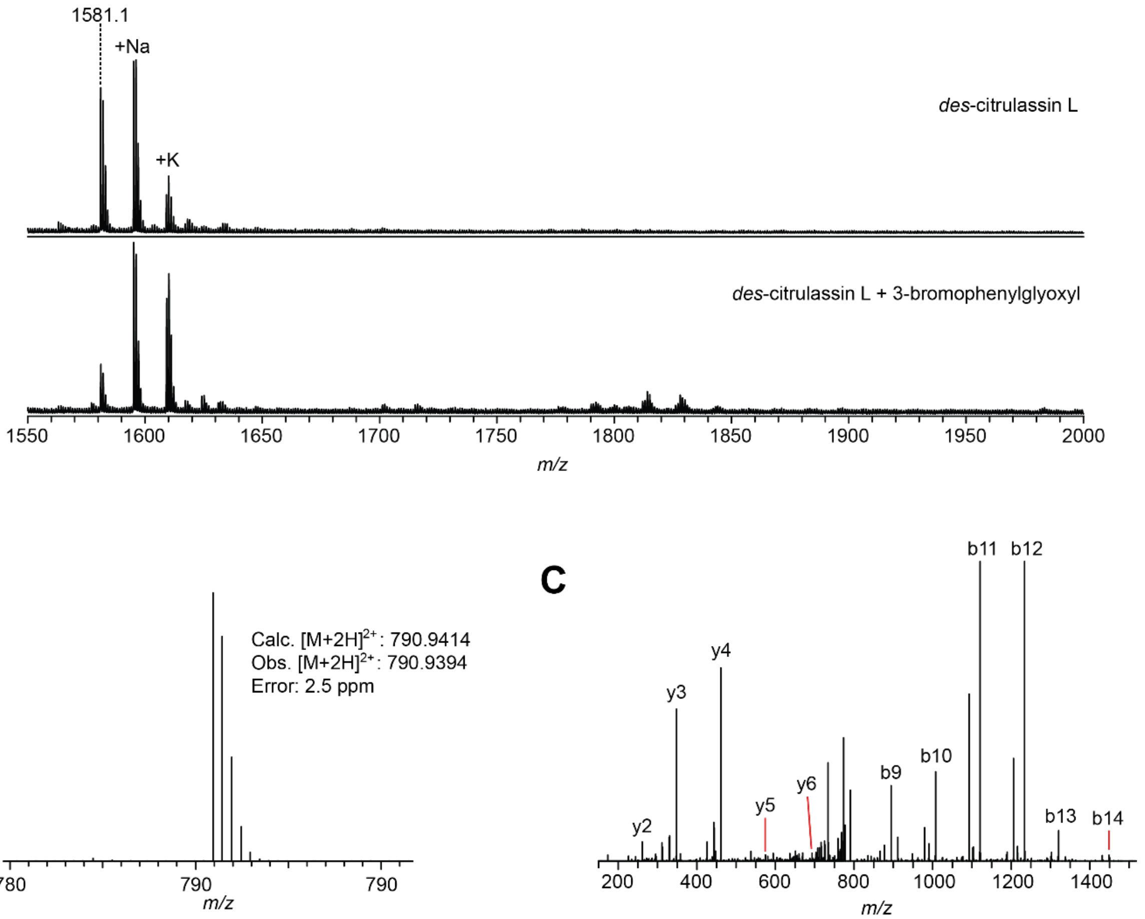

D

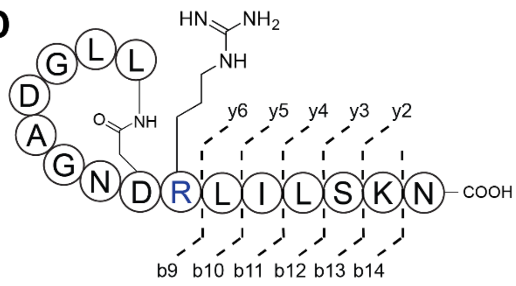

E

| Ion | Calc. mass | Obs. mass | Error (ppm) |
|---|---|---|---|
| **b14** | 1448.8220 | 1448.8176 | 3.0 |
| **b13** | 1320.7270 | 1320.7233 | 2.8 |
| **b12** | 1233.6950 | 1233.6913 | 3.0 |
| **b11** | 1120.6109 | 1120.6076 | 2.9 |
| **b10** | 1007.5269 | 1007.5238 | 3.1 |
| **b9** | 894.4428 | 894.4401 | 3.0 |
| **y2** | 261.1557 | 261.1552 | 2.0 |
| **y3** | 348.1878 | 348.1870 | 2.2 |
| **y4** | 461.2718 | 461.2707 | 2.4 |
| **y5** | 574.3559 | 574.3544 | 2.6 |
| **y6** | 687.4400 | 687.4383 | 2.4 |

**Part xii: *des*-citrulassin M, *Streptomyces pharetrae* B-24333, alternate MS media**

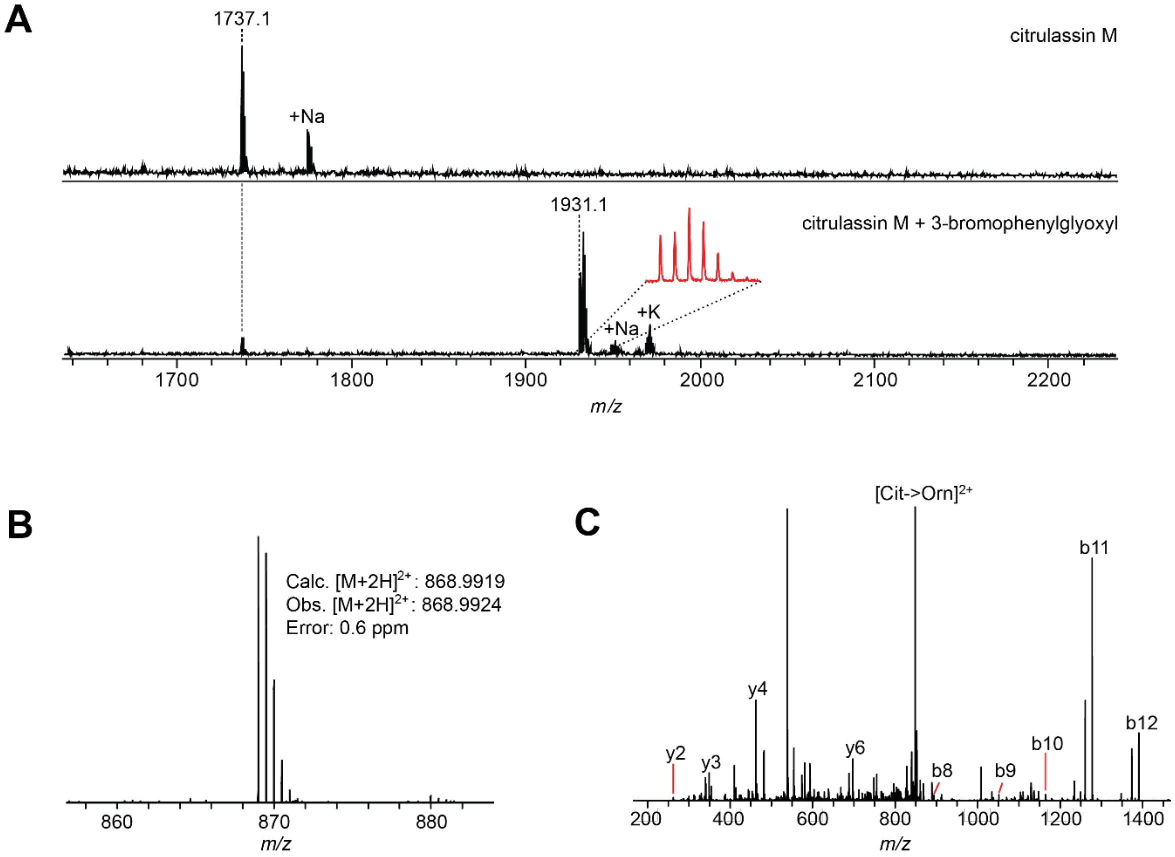

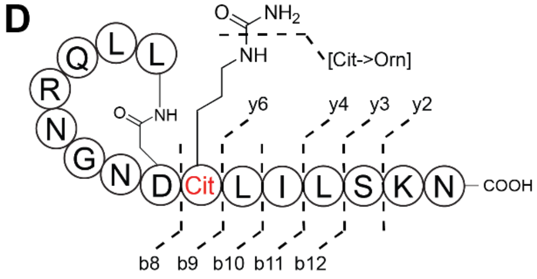

**E**

| Ion | Calc. mass | Obs. mass | Error (ppm) |
|---|---|---|---|
| **b12** | 1389.7967 | 1389.7965 | 0.1 |
| **b11** | 1276.7126 | 1276.7125 | 0.1 |
| **b10** | 1163.6285 | 1163.6302 | 1.5 |
| **b9** | 1050.5445 | 1050.5446 | 0.1 |
| **b8** | 893.4593 | 893.4595 | 0.2 |
| **y2** | 261.1557 | 261.1558 | 0.4 |
| **y3** | 348.1878 | 348.1879 | 0.3 |
| **y4** | 461.2718 | 461.2720 | 0.4 |
| **y6** | 687.4400 | 687.4407 | 1.0 |
| **[Cit-> Orn]$^{2+}$** | 847.4890 | 847.4893 | 0.4 |

**Table S3: PAD Pfam analysis.** PADs from *Homo sapiens*, (PAD1: NP_037490.2; PAD2: BAA82557.1; PAD3: NP_057317.2; PAD4: AAH25718.1; PAD6: NP_997304.3), *S. albulus* B-3066 (WP_064069847.1), and *Pseudomonas gingivalis* (WP_005873463.1) were analyzed against the pHMMs for PF03068 and PF04371 using hmmscan (https://www.ebi.ac.uk/Tools/hmmer/search/hmmscan). Members of PF03068 and PF04371 are shown in blue and orange, respectively.

| | **PF03068** | | **PF04371** | |
|---|---|---|---|---|
| NCBI Accession | Alignment Score | E-value | Alignment Score | E-value |
| **NP_037490.2** | 564 | 4.9E-173 | -1.1 | 0.34 |
| **BAA82557.1** | 553 | 1.3E-169 | 3.9 | 0.011 |
| **NP_057317.2** | 560 | 7.9E-172 | -0.5 | 0.24 |
| **AAH25718.1** | 573 | 1.1E-175 | 2.0 | 0.041 |
| **NP_997304.3** | 498 | 5.5E-153 | 2.2 | 0.036 |
| **WP_064069847.1** | 396 | 6.6E-122 | -0.3 | 0.21 |
| **WP_005873463.1** | 0.67 | -2.30 | 107 | 6.1E-34 |

**Table S4: Percent identity and percent similarity matrix of PADs.** PADs from *H. sapiens*, (PADI: NP_037490.2; PADII: BAA82557.1; PADIII: NP_057317.2; PADIV: AAH25718.1; PADVI: NP_997304.3) *S. albulus* B-3066 (WP_064069847.1), and *Pseudomonas gingivalis* (WP_005873463.1), the former of which are members of protein family PF03068 and the last of which is a member of PF04371. Because PF03068 and PF04371 PADs have different domain architectures, the excised PADIV (residues 301-663)[11] and the *P. gingivalis* PAD (residues 49-360)[12] catalytic domains were compared as well. The identity matrix was generated using EMBL-EBI Multiple Sequence comparison by Log-Expectation (MUSCLE)[13] and the Sequence Manipulation Suite.[14]

| | Species | *P. gingivalis* | *P. gingivalis* cat. dom. | *S. albulus* | *H. sapiens* PADVI | *H. sapiens* PADII | *H. sapiens* PADI | *H. sapiens* PADIII | *H. sapiens* PADIV | *H. sapiens* PADIV cat. dom. |
|---|---|---|---|---|---|---|---|---|---|---|
| **Species** | **Pfam** | PF04371 | PF04371 | PF03068 | PF03068 | PF03068 | PF03068 | PF03068 | PF03068 | PF03068 |
| ***P. gingivalis*** | PF04371 | 100 | 100 | 33 | 30 | 33 | 32 | 32 | 31 | 31 |
| ***P. gingivalis* cat. dom.** | PF04371 | 100 | 100 | 36 | 33 | 35 | 33 | 33 | 32 | 30 |
| ***S. albulus*** | PF03068 | 11 | 14 | 100 | 42 | 46 | 45 | 47 | 45 | 52 |
| ***H. sapiens* PADVI** | PF03068 | 11 | 13 | 23 | 100 | 61 | 61 | 62 | 61 | 64 |
| ***H. sapiens* PADII** | PF03068 | 13 | 13 | 26 | 44 | 100 | 64 | 66 | 63 | 72 |
| ***H. sapiens* PADI** | PF03068 | 13 | 14 | 26 | 46 | 53 | 100 | 69 | 69 | 80 |
| ***H. sapiens* PADIII** | PF03068 | 14 | 15 | 29 | 45 | 52 | 58 | 100 | 67 | 77 |
| ***H. sapiens* PADIV** | PF03068 | 13 | 14 | 26 | 46 | 51 | 58 | 56 | 100 | 100 |
| ***H. sapiens* PADIV cat. dom.** | PF03068 | 12 | 14 | 32 | 53 | 61 | 69 | 70 | 100 | 100 |

| % identity | % similarity |
|---|---|

**Figure S6: Protein sequence alignment of the *H. sapiens* and *S. albulus* PAD amino acid sequences.** Secondary structure features from the *H. sapiens* PAD (PDB code: 4DKT) are shown above the sequence alignment with the *S. albulus* PAD (WP_064069847.1). Identical residues are indicated by a red filled square, while similar residues are indicated by a white filled square (see legend below). Alignment was generated by MUSCLE with default settings and visualized by ESPRIPT3 (http://espript.ibcp.fr).[15]

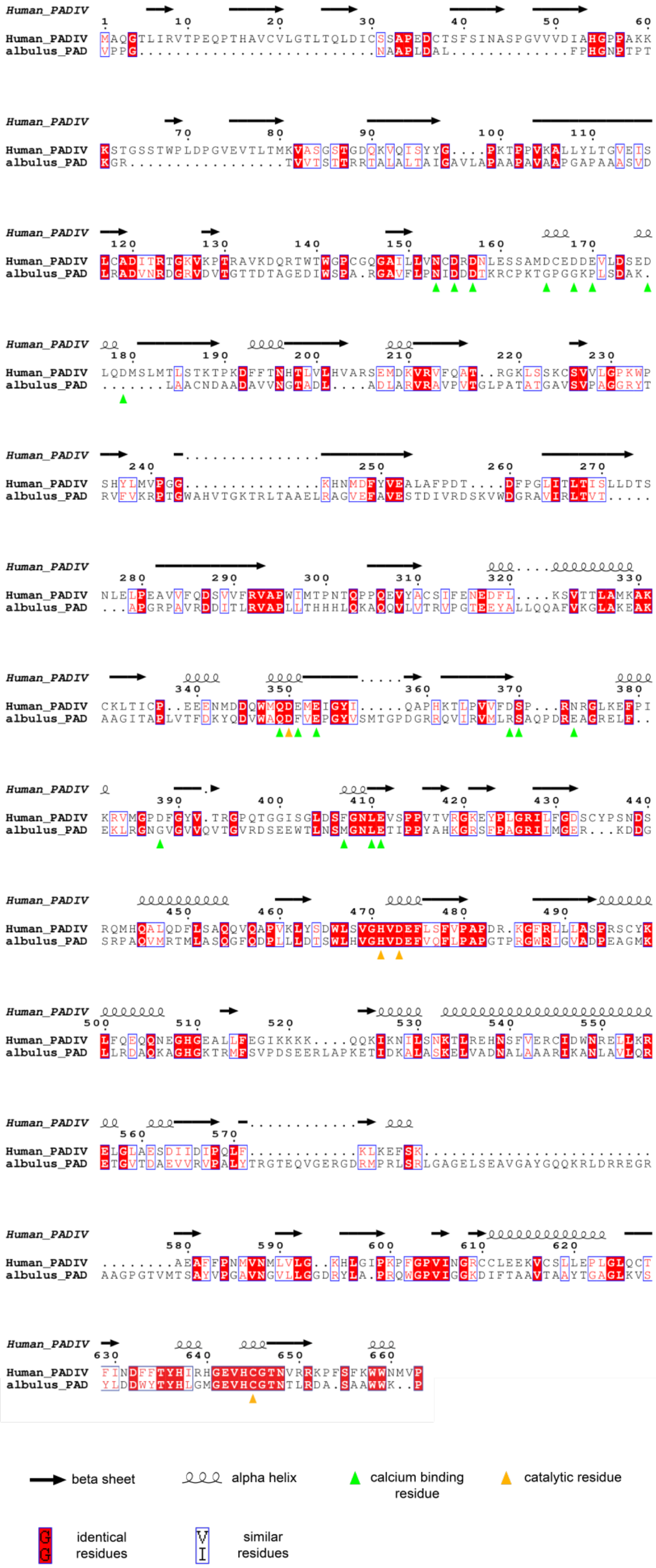

**Figure S7: Protein sequence alignment of the *H. sapiens* PAD, *S. albulus* PAD, and *P. gingivalis* PAD catalytic domain amino acid sequences.** Residues identical between *H. sapiens* (4DKT_A) PADIV, *S. albulus* PAD (WP_064069847.1) and the *P. gingivalis* PAD catalytic domain (WP_005873463.1, residues 49-360) are indicated by a filled red square, while similar residues are indicated by a filled white square. Residues that are identical between only the *S. albulus* PAD and one of the two other proteins are colored red (see legend below). Alignment was generated by MUSCLE with default settings and visualized by ESPRIPT3.

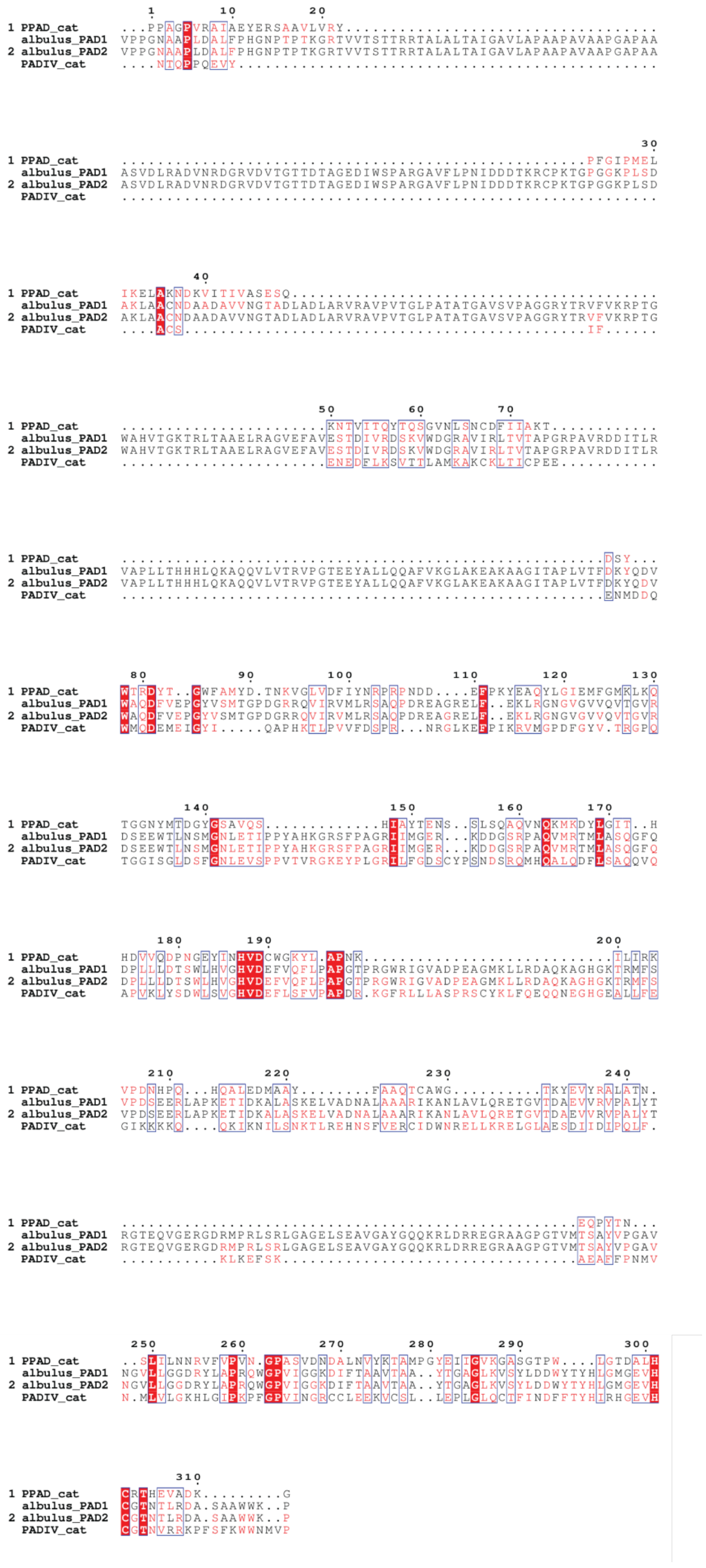

**Figure S8: Phylogenetic tree of bacterial PADs.** Circular phylogenetic tree of all PADs 500-800 amino acids in length, generated using standard settings on the Interactive Tree of Life (http://itol.embl.de/, see methods).[7] The outer ring is colored based on phylum of the PAD-containing organism (see legend) and is rooted based on PADIV (AAH25718.1).

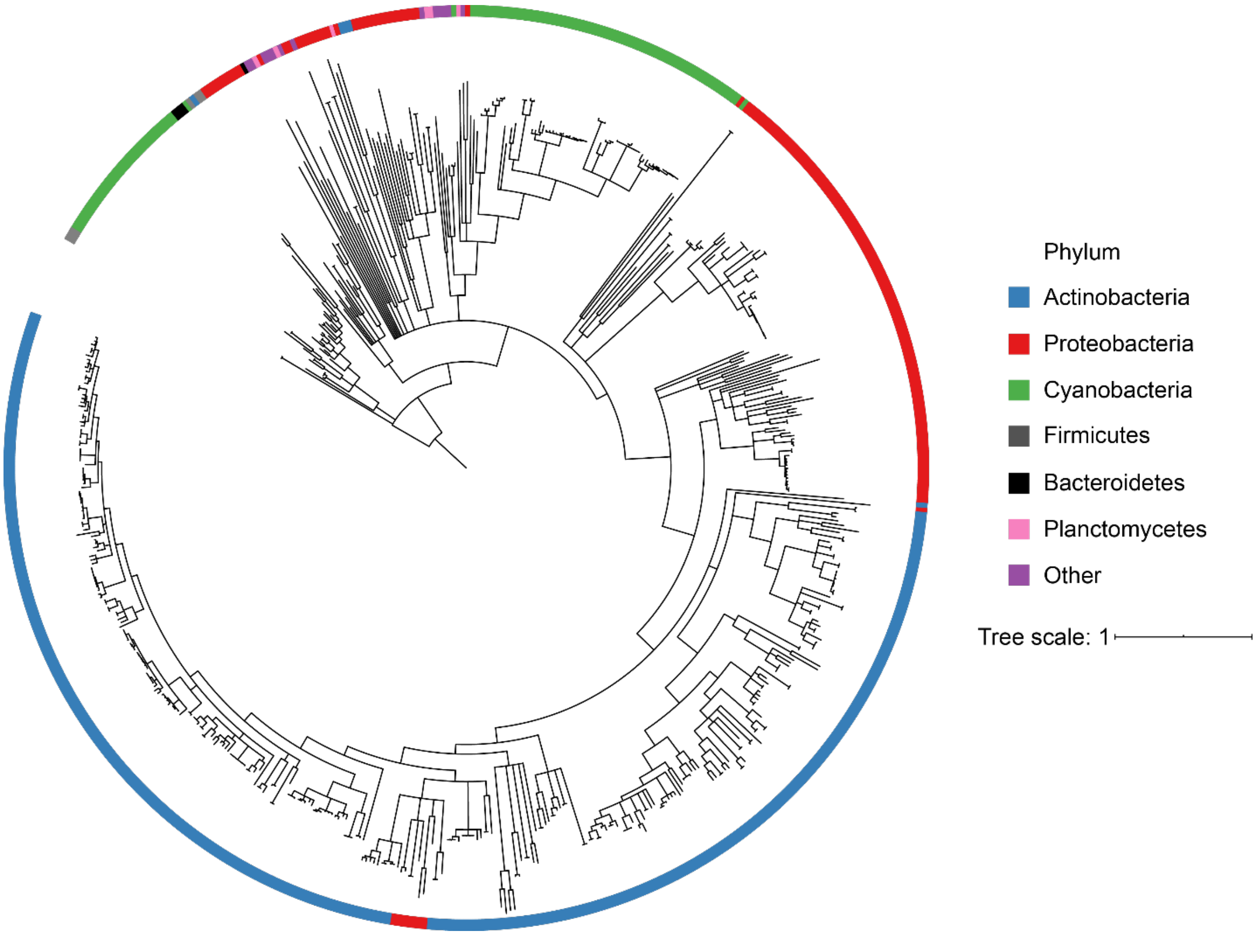

**Figure S9: PAD %GC.** (A) Parametric %GC analysis for horizontal gene transfer was performed by comparing the %GC of 672 bacterial PADs versus the organism genome. Linear regression was calculated and >2σ outliers are highlighted in blue. (B) Table of %GC content outliers, with the organism, NCBI protein accession identifier, PAD gene %GC, and organism genome %GC listed.

A

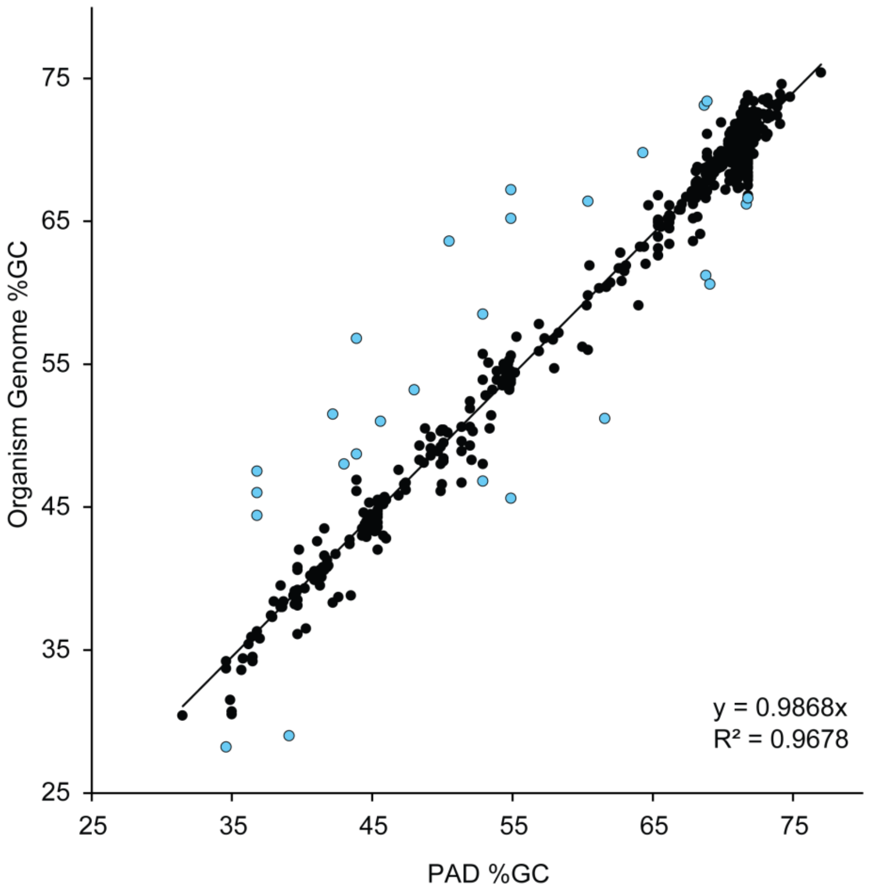

B

| Species | Protein accession | PAD%GC | Genome %GC | Difference |
|---|---|---|---|---|
| Bacterium isolate UCL2 | KAA0213793.1 | 51 | 64 | 13 |
| Methylocucumis oryzae | KJV04995.1 | 44 | 57 | 13 |
| Deltaproteobacteria bacterium isolate | TVQ88026.1 | 55 | 67 | 12 |
| Uncultured bacterium ACD_39C01209 | EKD82568.1 | 37 | 48 | 11 |
| Deltaproteobacteria bacterium isolate | RLB64542.1 | 55 | 65 | 10 |
| Candidatus Kentron sp. LPFa | VFK14916.1 | 42 | 52 | 10 |
| Uncultured bacterium ACD_39C00839 | EKD83174.1 | 37 | 46 | 9 |
| Candidatus Cloacimonetes bacterium isolate NORP72 | PCJ16673.1 | 39 | 29 | -10 |
| Sedimenticola sp. isolate BM503 | PLY16382.1 | 62 | 51 | -11 |
| Deltaproteobacteria bacterium isolate | RLC13998.1 | 55 | 46 | -9 |
| Uncultured bacterium ACD_47C00183 | EKD69285.1 | 37 | 44 | 7 |
| Pseudorhodoferax sp. Leaf265 | WP_056665489.1 | 69 | 61 | -8 |
| Planctomycetes bacterium isolate | RLS82094.1 | 60 | 66 | 6 |
| Rubrivivax sp. isolate PMG_229 | RZI79885.1 | 69 | 61 | -8 |
| Leptolyngbya sp. KIOST-1 | WP_035984404.1 | 53 | 59 | 6 |
| Myxococcales bacterium isolate | RYE93026.1 | 64 | 70 | 6 |
| Microcoleus sp. IPPAS B-353 | WP_159789547.1 | 46 | 51 | 5 |
| Lachnotalea glycerini | OYO86896.1 | 35 | 28 | -7 |
| Oscillatoria sp. PCC 10802 | WP_082218359.1 | 48 | 53 | 5 |
| Chitinophagaceae bacterium isolate PMG_220 | RYY52335.1 | 43 | 48 | 5 |
| Methylocucumis oryzae | KJV05480.1 | 44 | 49 | 5 |
| Leptolyngbya sp. PCC 7375 | WP_006513830.1 | 53 | 47 | -6 |
| Myxococcus sp. AM301 | WP_163782873.1 | 69 | 73 | 4 |
| Nitriliruptoraceae bacterium ZYF776 | WP_155383327.1 | 69 | 73 | 4 |
| Streptomyces halstedii | WP_164347388.1 | 72 | 66 | -6 |
| Streptomyces sp. SID13031 | WP_164601292.1 | 72 | 67 | -5 |

**Table S5: Local Co-occurrence analysis for bacterial PADs.** Protein Families (Pfams) with ≥10% co-occurrence are shown below. A comprehensive list can be found in the Supplementary Dataset. Bolded are PAD PFAMs.

| Pfam | Count | Percent | Title | Description |
|---|---|---|---|---|
| **PF03068** | **686** | **102** | **PAD** | **protein-arginine deiminase** |
| PF00005 | 201 | 30 | ABC_trans | ABC transporter transmembrane region |
| **PF08527** | **189** | **28** | **PAD_M** | **protein-arginine deiminase middle domain** |
| PF13304 | 170 | 25 | AAA_21 | AAA domain, putative AbiEii toxin, type IV TA system |
| PF00072 | 155 | 23 | Response_reg | Response regulator receiver domain |
| PF07690 | 135 | 20 | MFS_1 | major facilitator superfamily |
| PF00440 | 134 | 20 | TetR_N | bacterial regulatory proteins, tetR family |
| PF00583 | 116 | 17 | Acetyltransf_1 | Acetyltransferase (GNAT) family |
| PF00528 | 96 | 14 | BPD_transp_1 | binding-protein-dependent transport system inner membrane component |
| PF00196 | 94 | 14 | GerE | bacterial regulatory proteins, luxR family |
| PF02463 | 90 | 13 | SMC_N | RecF/RecN/SMC N terminal domain |
| PF02518 | 87 | 13 | HATPase_c | histidine kinase-, DNA gyrase B-, and HSP90 like ATPase |
| PF13673 | 86 | 13 | Acetyltransf_10 | Acetyltransferase (GNAT) domain |
| PF00561 | 85 | 13 | Abhydrolase_1 | alpha/beta hydrolase fold |
| PF08281 | 81 | 12 | DUF1492 | protein of unknown function (DUF1492) |
| PF13508 | 78 | 12 | Acetyltransf_7 | Acetylatransferase (GNAT) domain |
| PF00106 | 75 | 11 | adh_short | short chain dehydrogenase |
| PF12802 | 74 | 11 | MarR_2 | MarR family |
| PF13560 | 74 | 11 | HTH_31 | Helix-turn-helix domain |
| PF01381 | 71 | 11 | HTH_3 | Helix-turn-helix |
| PF13561 | 70 | 10 | adh_short_C2 | enoyl-(Acyl carrier protein) reductase |
| PF07719 | 68 | 10 | TPR_2 | Tetracopeptide repeat |
| PF08659 | 68 | 10 | KR | KR domain |
| PF12697 | 67 | 10 | Abhydrolase_6 | alpha/beta hydrolase family |

**Figure S10: Local genomic neighborhood of bacterial PADs.** Shown are nine examples of bacterial PADs with four predicted coding sequences on each side of the PAD-encoding gene displayed. These PADs (blue ORFs) do not appear in a conserved genomic context. Pfam descriptions are described on the right for each neighboring gene.

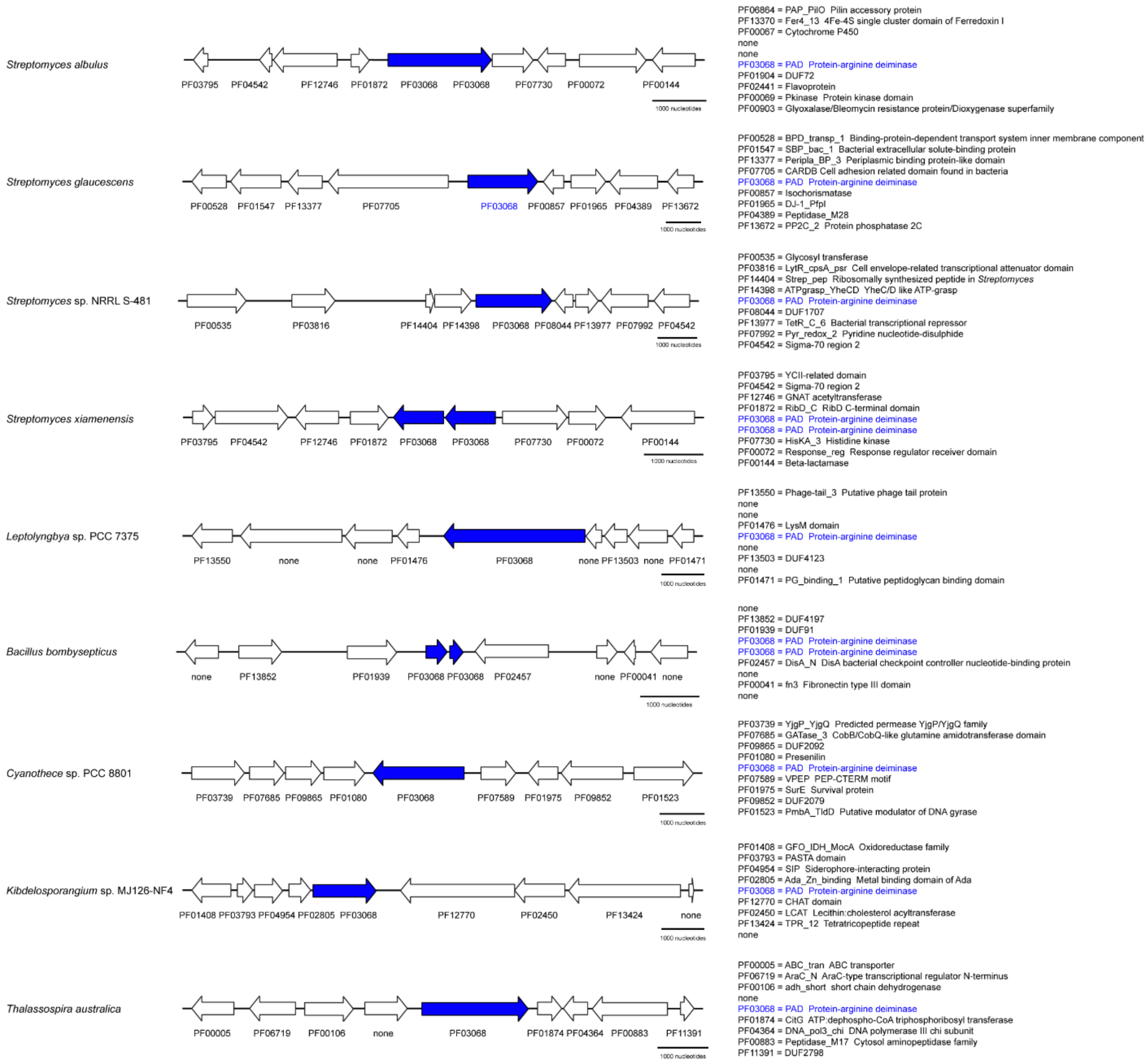

**Figure S11: PCR confirmation of chromosomal PAD insertion.** *ermE*p*-PAD insert from the conjugative pAE4 plasmid used in this study is compared with gDNA PCR products from wild-type *S. katrae* B-16271 and randomly selected pAE4 exconjugates. PCR fragment containing *ermE*p*-PAD is ~2.7 kb.

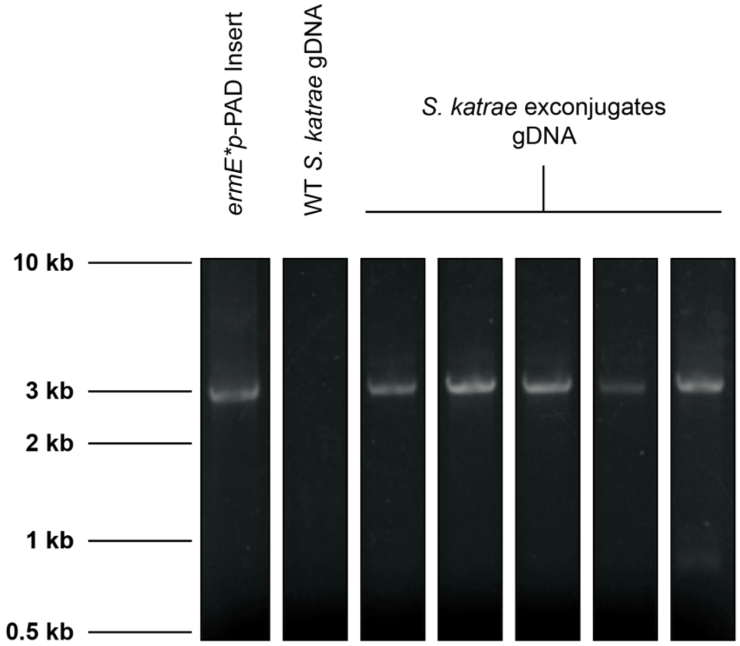
